# Tethering ferredoxin-NADP^+^ reductase to photosystem I promotes photosynthetic cyclic electron transfer

**DOI:** 10.1101/2024.11.01.621516

**Authors:** Thomas Z. Emrich-Mills, Matthew S. Proctor, Gustaf E. Degen, Philip J. Jackson, Katherine H. Richardson, Frederick R. Hawkings, Felix Buchert, Andrew Hitchcock, C. Neil Hunter, Luke C.M. Mackinder, Michael Hippler, Matthew P. Johnson

## Abstract

Fixing CO_2_ via photosynthesis requires ATP and NADPH. Linear electron transfer (LET) supplies both metabolites, yet depending on environmental conditions, additional ATP is required which can be generated by cyclic electron transfer (CET). How the balance between LET and CET is set remains largely unknown. Ferredoxin(FD)-NADP^+^ reductase (FNR) has been suggested to act as the switch, channelling photosynthetic electrons to LET when it is bound to photosystem I (PSI) or CET when bound to cytochrome *b*_6_*f*. The essential role of FNR in LET precludes the use of a direct gene knock-out to test this hypothesis. We circumvented this problem by using CRISPR-Cas9 gene editing in *Chlamydomonas reinhardtii* to create a chimeric form of FNR tethered to PSI via PSAF. Chimeric FNR mutants exhibited impaired photosynthetic growth relative to the wild type, and decreased LET and non-photochemical quenching (NPQ) under high CO_2_ conditions. However, higher rates of CET and increased NPQ were found in the mutants under low CO_2_ or low O_2_ conditions where CET is normally up-regulated. Thus, rather than PSI-FNR interactions promoting LET, tethering FNR to PSI promotes CET at the expense of LET and CO_2_ fixation.

## Introduction

Photosynthesis is essential to life on Earth, converting solar energy into biomass which sustains global food chains, while generating oxygen as a by-product. The first steps of oxygenic photosynthesis take place in the chloroplast thylakoid membrane. Here specialised light harvesting complexes (LHCs) absorb solar energy and transfer the excitation to the photosystem I and II (PSI, PSII) reaction centres, where photochemistry takes place (Nelson and Junge 2015). PSII uses the excitation energy to split water into protons and electrons and reduce plastoquinone (PQ) to plastoquinol (PQH_2_). PQH_2_ is then oxidised by the cytochrome *b*_6_*f* (cyt*b*_6_*f*) complex which reduces plastocyanin (PC) in the thylakoid lumen (Malone et al. 2021). The protons liberated by cyt*b*_6_*f* and PSII contribute to the proton motive force (pmf) across the thylakoid membrane, which is utilised by ATP synthase to power the endergonic formation of ATP (Kühlbrandt 2019). PSI uses excitation energy to oxidise PC and reduce ferredoxin (FD). Electrons from FD are utilised in linear electron transfer (LET) by the enzyme FD-NADP^+^ reductase (FNR) to reduce NADP^+^ to NADPH, which in turn is used to reduce 3-phosphoglycerate during the CO_2_-fixing Calvin-Benson-Bassham (CBB) cycle. Alternatively electrons from FD or NADPH can re-enter the photosynthetic electron transfer chain to reduce PQ to PQH_2_, via cyclic electron transfer (CET) (Yamori and Shikanai 2015; Nawrocki et al. 2019b). Since LET produces both ATP and NADPH, while CET produces only ATP, their relative activities must be carefully regulated to meet the varying energetic demands of CO_2_ fixation and other metabolic pathways in the chloroplast (Alric and Johnson 2017; Burlacot 2023). In the model eukaryotic green alga *Chlamydomonas reinhardtii*, several other alternative electron transfer pathways also contribute to the chloroplast pmf and so the ATP/ NADPH balance. These include the pseudo-cyclic electron transfer (PCET) via the flavodiiron proteins (FLVA and FLVB) which use FD to reduce O_2_ to water (Chaux et al. 2017). Alternatively, reductant may be exported from the chloroplast and consumed in the mitochondria via respiratory electron transfer (RET) (Cardol et al. 2009; Dang et al. 2014). The ATP produced via RET can then be imported into the chloroplast possibly via the triose phosphate translocator (Alric and Johnson 2017). Since each of these pathways make varying contributions to the pmf and ATP/NADPH balance depending on environmental conditions (Burlacot et al. 2022), their careful redox-dependent regulation is crucial to modulation of ΔpH-dependent photoprotection of PSII, via non-photochemical quenching (NPQ) of excess excitation energy, and of PSI, via photosynthetic control (PCON) (Murchie and Ruban 2020; Degen and Johnson 2024).

In Chlamydomonas, two mechanisms of CET have been shown to exist, one mediated by the non-proton-pumping NDA2 complex, which is NADPH-dependent (Desplats et al. 2009), and one that is FD-dependent (Alric et al. 2010; Alric 2014). A leading contender for the role of the FD-dependent plastoquinone reductase (FQR) was the complex formed between the Proton Gradient Regulation like 1 (PGRL1) and Proton Gradient Regulation 5 (PGR5) proteins (Munekage et al. 2004; DalCorso et al. 2008; Tolleter et al. 2011; Johnson et al. 2014). However, it was later shown that PGRL1 is not essential for CET in either Chlamydomonas or Arabidopsis (Nawrocki et al. 2019a; Rühle et al. 2021). Moreover, the proposed thiol-based interaction between PGRL1 and PGR5 was also not required for functional CET (Buchert et al. 2020). Enhanced CET activity can be measured in Chlamydomonas under low CO_2_, which increases ATP demand to fuel the carbon concentrating mechanism (CCM), and under low oxygen conditions (anoxia), which suppress PCET and RET (Lucker and Kramer 2013; Clowez et al. 2015; Burlacot et al. 2022). Anoxia and low CO_2_ promote reduction of NADP^+^ thus lowering the redox potential of the chloroplast stroma and triggering reduction of the PQ pool (Johnson and Alric 2012). In Chlamydomonas PQ reduction promotes the phosphorylation of PSII light harvesting antenna proteins (LHCII) by the kinase STT7, which detach from PSII and become associated with PSI (State II) (Depège et al. 2003; Nawrocki et al. 2015). The reverse transition to State I is promoted by oxidation of the PQ pool and is catalysed by the PPH1 and PBCP phosphatases (Cariti et al. 2020). Biochemical analysis correlated anoxia with the appearance of a high molecular weight supercomplex containing not only PSI and LHCII but also cyt*b*_6_*f*, its weakly bound auxiliary subunit PETO, Anaerobic Response 1 (ANR1), Calcium Sensor Kinase (CAS), PGRL1 and FNR (Iwai et al. 2010; Terashima et al. 2012; Takahashi et al. 2013). If NADPH and FD were added to this supercomplex preparation the *b*-type hemes associated with the PQ reduction site in cyt*b*_6_*f* were efficiently reduced, suggesting cyt*b*_6_*f*-tethered FNR mediates FQR activity (Iwai et al. 2010). Recently, it was shown that under anoxic conditions the oxidation of the *b*-type hemes in cyt*b*_6_*f* is accelerated in a PGR5-dependent fashion (Buchert et al. 2020). Since PGR5 affects the tethering of FNR to the thylakoid membrane it was suggested to modulate CET activity of cyt*b*_6_*f* (Mosebach et al. 2017). In this model the location of FNR therefore decides the fate of electrons emanating from PSI, with soluble or PSI bound FNR mediating LET, while cyt*b*_6_*f* bound FNR mediates CET (Joliot and Johnson 2011). In Arabidopsis, FNR tethering to the thylakoid membrane via TROL and TIC62 complexes was shown to promote CET, however these proteins are absent from Chlamydomonas (Kramer et al. 2021). Moreover, since the putative CET supercomplex is extremely fragile, to date only a low-resolution structure exists and no information on the precise interactions between FNR, PSI and cyt*b*_6_*f* exist to clarify the mechanism (Buchert et al. 2018; Steinbeck et al. 2018). Indeed, controversy remains since isolated FNR has itself been shown to possess FQR activity (Bojko et al. 2003), while it has also been suggested that FD could donate electrons directly to cyt*b*_6_*f* in the absence of FNR (Nawrocki et al. 2019b). Therefore, new approaches are required to resolve the outstanding questions regarding the role of FNR location and membrane binding on the balance between LET and CET in Chlamydomonas.

Unfortunately, due to the essential nature of FNR for both autotrophic and heterotrophic (e.g. in nitrogen and fatty acid metabolism (Terauchi et al. 2009)) growth in Chlamydomonas it has not been possible to test whether its interaction with cyt*b*_6_*f* is a prerequisite for CET. To address this issue, we produced a chimeric *FNR-PSAF* gene encoding a fusion protein that would ensure tethering of FNR to PSI, while simultaneously disrupting the native *FNR* and *PSAF* genes generating a Chlamydomonas mutant where the only FNR present is the chimeric protein. Unexpectedly, the PSAF-FNR chimeric mutants show enhanced CET compared to the wild-type UVM4 (WT) and restricted LET under high CO_2_ conditions. Our results therefore confirm the key role of FNR location in determining the fate of photosynthetic electrons between CET and LET but exclude the necessity for FNR binding to cyt*b*_6_*f* in the mechanism of CET.

## Results

### Creation of a chimeric PSAF-FNR protein in Chlamydomonas

To achieve our goal of selectively tethering FNR to PSI we began by designing a chimeric protein where FNR is fused to the stromal facing C-terminus of the PSI subunit PSAF using a small 9 amino acid linker sequence (Fig. 1A). The plasmid also contains an *AphVIII* selection marker, allowing selection of transformants on paromomycin (Li et al. 2016). The gene encoding the PSAF-FNR chimera was then introduced via CRISPR-Cas9 gene editing to the WT native nuclear *PSAF* locus disrupting the gene. The independent mutant lines created (NT4 and NT12), contained both native *FNR* and chimeric *PSAF-FNR* genes (Fig. 1B) and the corresponding proteins (Figs. 1C and 1D). The NT4 and NT12 mutant lines were then further modified using CRISPR-Cas9 to inactivate the native *FNR* gene by inserting an *AphVII* hygromycin resistance cassette (Fig. 1E), producing stable independent lines (T7 and T10). Immunoblotting with anti-PSAF and anti-FNR antibodies confirmed that the T7 and T10 mutant cell lines lacked the native PSAF and FNR proteins and contained only the PSAF-FNR chimeric protein at ∼53 kDa (Figs. 1F and 1G). The whole cell immunoblots also failed to detect any evidence that the chimeric PSAF-FNR protein was proteolytically cleaved in the mutants to yield free FNR or PSAF.

**Figure 1.**
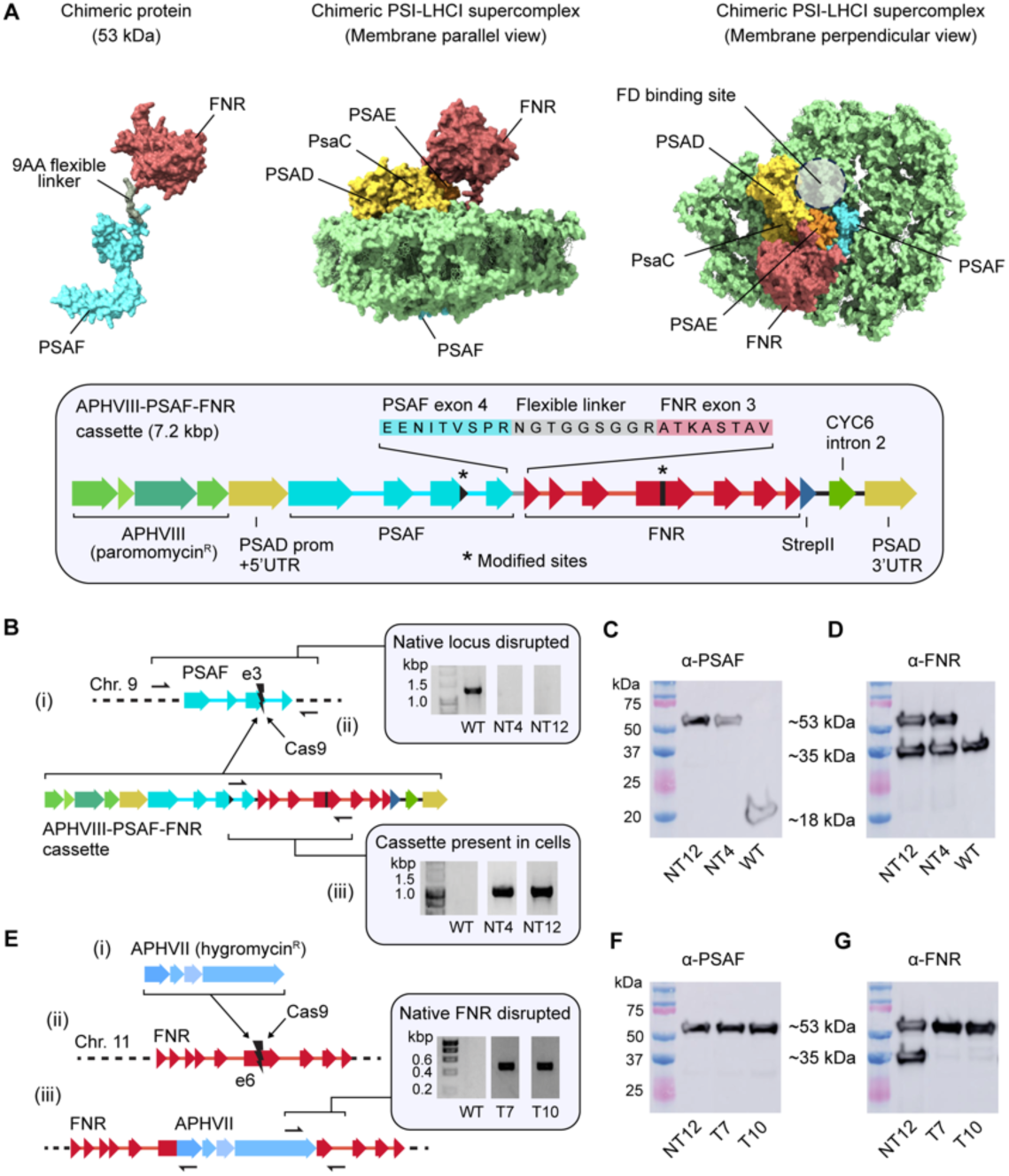
| Creation of chimeric PSAF-FNR mutants in Chlamydomonas. **A**, Putative molecular model of the PSAF-FNR chimeric PSI-LHCI complex. The inset below shows a map of the expression cassette for the chimeric protein, along with the amino acid sequence of the linker region. Modified sites refer to silent mutations made in *PSAF* exon 3 and *FNR* exon 6 to avoid CRISPR targeting. **B**, Results of the co-transformation of WT with the *PSAF-FNR* expression cassette and Cas9-single guide RNA ribonucleoprotein (RNP) targeting *PSAF* exon 3. (i) shows the intended outcome of CRISPR-guided insertional mutagenesis, in which the *PSAF-FNR* expression cassette is inserted into native *PSAF* exon 3 as part of the NHEJ repair process. (ii) shows colony PCRs with transformant lines NT4 and NT12. Absence of an amplicon across the *PSAF* locus indicated a large insertion at exon 3; the black arrows indicate the priming sites used to detect this amplicon by PCR. (iii) shows successful amplification using primers specific to the *PSAF-FNR* cassette in lines NT4 and NT12, which indicated successful integration of the cassette into the genome. **C**, Immunoblots with anti-PSAF antibody confirmed deletion of native PSAF in NT4 and NT12. **D**, immunoblots with an anti-FNR antibody confirmed expression of the PSAF-FNR protein along with native FNR. **E**, Results of the CRISPR-mediated deletion of native *FNR*. (i) shows the *AphVII* cassette used for transformation, encoding resistance to hygromycin. The intention was to select for insertion of *AphVII* into native *FNR* exon 6 (ii) as part of the NHEJ repair process. (iii) shows the intended result, along with evidence of successful PCR amplification across the boundary between native *FNR* and *AphVII* using a three-primer PCR that allowed for insertion of *AphVII* in either orientation. Amplicons in transformant lines T7 and T10 are strong evidence of correct insertion. **F**, T7 and T10 immunoblots using anti-PSAF and **G**, anti-FNR antibodies indicate loss of native FNR and the presence of the PSAF-FNR chimera.

### Chlamydomonas mutants expressing the *PSAF-FNR* chimeric gene show inhibited photoautotrophic growth

We next assessed the photoautotrophic growth of the mutants versus the WT at a light intensity of 60 μmol photons m^-2^ s^-1^ using 14 h/10 h light-dark cycle and atmospheric CO_2_ as the sole carbon source (TP medium) (Fig 2A). The T7, T10, NT4 and NT12 mutants all showed increased doubling time compared to the WT (Fig 2A and A). In contrast, photomixotrophic growth under the same light conditions while employing acetate as the carbon source was unaffected (TAP medium) (Fig 2C). This suggests that the lesion caused by the introduction of the chimera affects photosynthetic electron transfer in such a way as to impede the efficient assimilation of CO_2_. In line with this, we recorded a lower rate of PSII electron transfer (ETRII) in the mutants compared to the WT in photoautotrophically grown cultures, even when cells were supplied with a saturating amount (10 mM) of sodium bicarbonate (Fig 2D).

**Figure 2.**
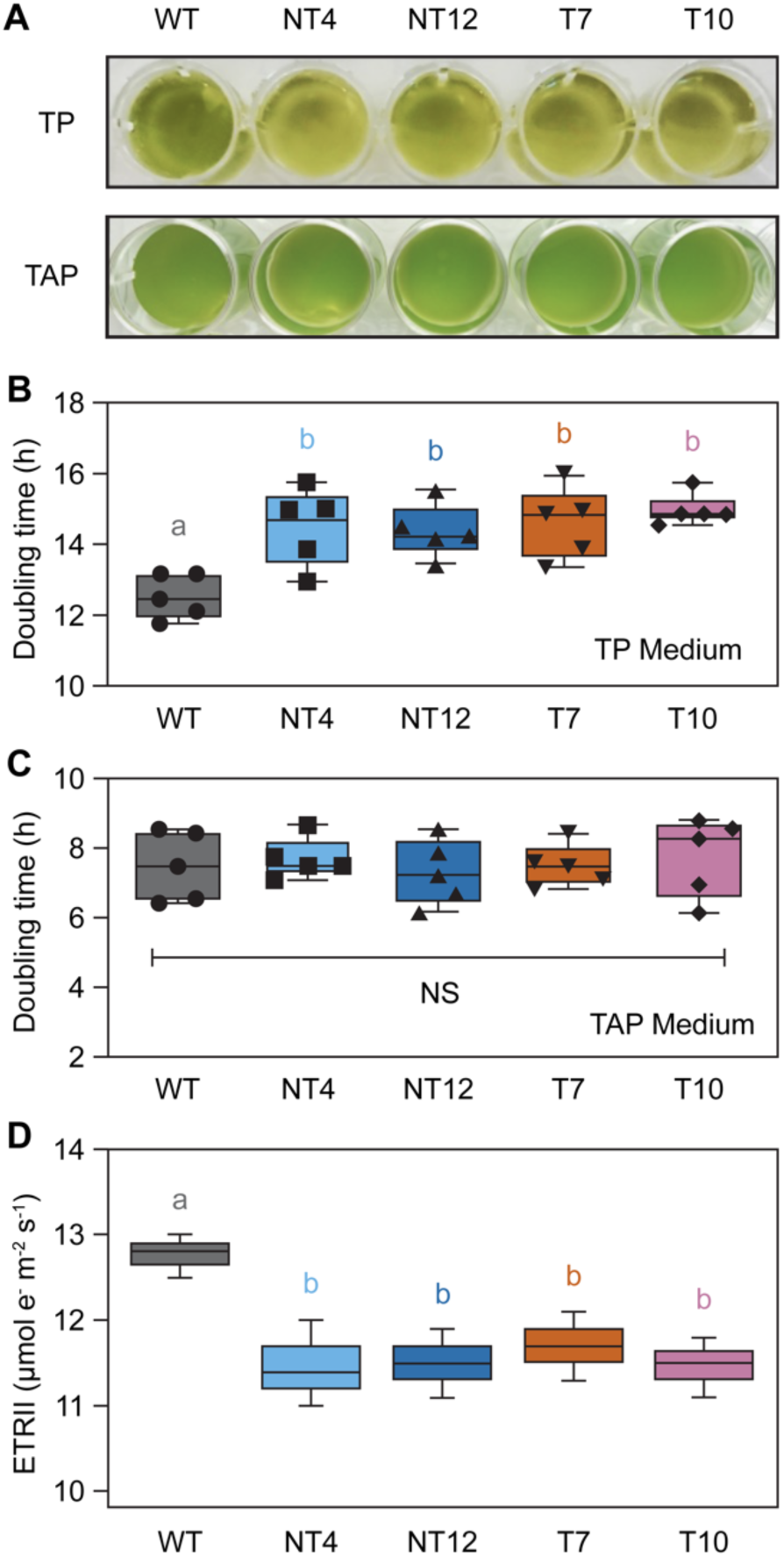
| Growth and electron transport phenotype of chimeric PSAF-FNR mutants. A, Representative image of photoautotrophic (TP medium) and photomixotrophic (TAP medium) growth for the chimeric PSAF-FNR mutants. **B**, Average doubling time in hours for growth in TP medium. **C**, Average doubling time in hours for growth in TAP medium. For A-C the growth conditions used were ∼60 μmol photons m^-2^ s^-1^ (white LED) 14 h/ 10 h day-night cycling, shaking at 120 rpm at 20 °C. Error bars are S.E.M. (three replicates per strain). **D**, PSII electron transport rate (ETRII) determined via chlorophyll fluorescence for the WT and PSAF-FNR mutants at ∼60 μmol photons m^-2^ s^-1^ in the presence of 10 mM sodium bicarbonate. Median values are plotted. Error bars show the range (*n* = 3). Means were compared between genotypes using an ordinary two-way ANOVA and corrected using Tukey’s multiple comparison test followed by a HSD test with alpha = 0.05. Different letters indicate significant differences between data points.

### Mutant cells with the chimeric PSAF-FNR protein are shifted to State II

We selected the T7 and NT12 mutants for more detailed phenotypic analysis compared to the WT. First, we confirmed that the chimeric PSAF-FNR protein was stably incorporated into PSI by purifying thylakoids from the mutants, solubilising them with the detergent lauryl maltose neopentyl glycol (LMNG) and separating the complexes via sucrose gradient ultracentrifugation (Fig. 3A). The WT showed four major bands on the gradients, consistent with past studies, belonging to monomeric LHCII (band 1), trimeric LHCII (band 2), PSII (band 3) and PSI (band 4) (Fig. 3A) (Iwai et al. 2010; Takahashi et al. 2013; Buchert et al. 2018). An additional green band (band 5) was present in the gradients of both the NT12 and T7 mutants and band 4 was also of lower intensity compared to the WT (Fig. 3A). Immunoblotting with an anti-PSAE antibody revealed the presence of PSI in band 4 and in the mutants additionally in band 5 (Fig. 3B). The anti-FNR antibody confirmed a wide distribution of FNR throughout the gradient in the WT (fractions 2-11), consistent with previous results (Mosebach et al. 2017; Buchert et al. 2018). In contrast in NT12, both native ∼35 kDa and chimeric (∼53 kDa) proteins were detected and were concentrated predominantly in fractions 7-11, with a peak in intensity similar to PSI (Fig. 3B). In the T7 mutant, only the 53 kDa chimeric protein was detected and was again largely absent from fractions 1-5, coinciding with PSI distribution (Fig. 3B). The position of band 5 is consistent with that previously reported for the PSI-LHCII-cyt*b*_6_*f* supercomplex often observed in State II in Chlamydomonas (Iwai et al. 2010; Terashima et al. 2012; Takahashi et al. 2013). However, immunoblotting for the Rieske iron-sulphur protein subunit of cyt*b*_6_*f* (PETC) revealed this complex was largely absent from band 5 (fraction 8) and instead concentrated in fractions 4-6 in both mutants (Fig. 3B). We repeated the immunoblotting for fraction 8 using anti-PETC and anti-PetA antibodies with a longer exposure compared to Fig. 3B and this uncovered the presence of cyt*b*_6_*f* in band 5 (Fig. 3C), though this was poorly correlated with PSI as revealed by anti-PSAE antibody, with relatively higher cyt*b*_6_*f*/PSI ratio in the WT compared to the mutants. In contrast, the absorption spectrum of band 5 in the mutants revealed the clear presence of additional chlorophyll *b* in the 470 and 650 nm regions consistent with the presence of LHCII complexes (Fig. 3D). Enhanced fluorescence emission from PSI at 720 nm relative to PSII emission at 685 nm (Fig. 3E) showed that NT12 and T7 cells were in State II (Fig. 3E). The increased presence of LHCII, revealed by immunoblotting with the anti-LHCBM5 antibody, in fractions 7 and 8 of the NT12 and T7 mutants compared to the WT supported this assignment (Fig. 3B). Interestingly, the mutant cells adopted State II more readily than the WT under four different conditions that paired high or low CO_2_ with oxic or anoxic conditions (Supplementary Fig. S1A). However, the total capacity to switch to State II was similar in the WT and mutants, as verified by treating cells with nigericin which collapses ΔpH and drives cells fully to State II (Bulté et al. 1990) (Supplementary Fig. S1B). The chimeric protein therefore successfully associates with PSI, causes a redistribution of native FNR in NT12 and also results in a shift from State I to State II in both mutants.

**Figure 3.**
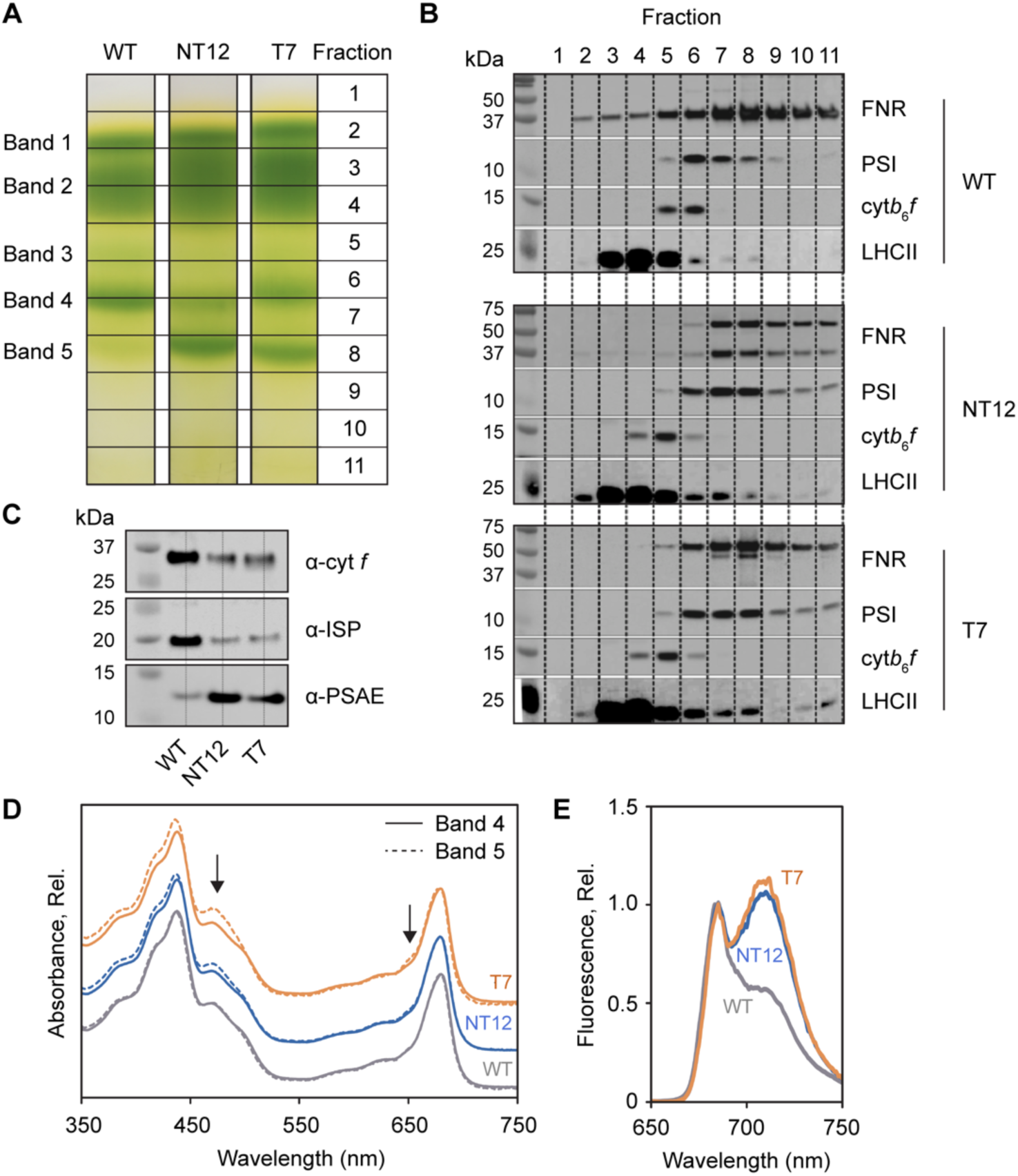
| Localisation of the PSAF-FNR chimera within PSI. A, Sucrose density gradients of LMNG solubilised thylakoids from WT, NT12 and T7 cells. Principle bands and fractions are numbered. **B**, Immunoblots (based on equal volume loading by sucrose gradient fraction) using anti-FNR, anti-PSAE (PSI), anti-PETC (cyt*b*_6_*f*) and anti-LHCBM5 (LHCII) antibodies. **C**, Immunoblots (based on equal volume loading of sucrose gradient fraction 8 alone) using anti-PetA, anti-PETC and anti-PSAE antibodies. **D**, Absorption spectra of fractions 6 and 8 from WT, NT12 and T7, normalised to the maximum intensity at 678 nm. **E**, 77K fluorescence emission spectra of WT, NT12 and T7 cells grown in TP at 60 μmol photons m^-2^ s^-1^ 14 h/ 10 h day-night cycling, shaking at 120 rpm.

### Chimeric PSAF-FNR mutants show increased PSII and decreased PSI abundance

To assess effects on the relative abundance of key photosynthetic proteins, we performed a proteomic analysis of WT and mutant cells via label-free quantitative (LFQ) mass spectrometry (MS), as previously described (Flannery et al. 2021). Proteins were extracted from whole cell lysates produced from four independent (biological) replicate cultures for each of the WT, NT12 and T7 lines grown under 60 μmol photons m^-2^ s^-1^ in TP medium. The biological replicates were analysed with three technical repeats. The 36 MS datafiles were then processed for protein identification and quantification using MaxQuant (Cox and Mann 2008) to generate iBAQ abundance scores (Schwanhäusser et al. 2011) to reveal differential protein levels. Using this approach, we first normalised each iBAQ score to the intra-analysis sum of the subunit iBAQ scores from proteins of the core photosynthetic machinery: PSI, PSII, cyt*b*_6_*f* and ATP synthase, listed in Source data. Using the median of three technical repeats to represent each biological replicate, we then applied a modified *t* test incorporating permutation-based false discovery rate with 250 randomisations, as implemented in Perseus, to indicate significant differences in the abundance of selected proteins in WT vs. mutant cells. The total number of proteins identified across all 36 analyses was 4091 (Source data). Fig. 4A shows FNR and PSAF levels calculated from their individual, summed tryptic peptide ion intensities since iBAQ would be inappropriate for separate quantification of PSAF and FNR in chimeric PSAF-FNR. NT12-expressed FNR, in both native and chimeric forms combined, was produced at 3.6-fold greater abundance relative to WT. In T7, expressing only chimeric PSAF-FNR, the level of FNR was 2.4-fold greater than WT. The difference in FNR level between NT12 and T7 is consistent with the additional contribution of native FNR in NT12 to the total FNR abundance against the background of enhanced FNR expression, as PSAF-FNR, in both mutants. PSAF levels, expressed only as PSAF-FNR in the mutants, were increased, though the difference was not statistically significant (Fig. 4A). In the NT12 and T7 mutants, the abundance of PSII was increased to 1.1-1.2 fold relative to the WT, while the abundance of PSI decreased to 0.74-0.68 fold (Fig. 4B). The decrease in PSI/PSII ratio in the mutants was confirmed by comparing the electrochromic shift (ECS) signal upon a single turnover flash in the presence and absence of 3-(3,4-dichlorophenyl)-1,1-dimethylurea (DCMU) and hydroxylamine (HA) to inhibit PSII (Supplementary Fig. S2A). On the other hand, there was no significant difference in the relative abundance of PC, cyt*b*_6_*f* and ATP synthase complexes in both mutants (Fig. 4B). The abundance of individual PSI subunits in the mutants followed a similar pattern to that of the whole complex for PsaA and PsaB, though for PsaC, PSAG, PSAH, PSAK and PSAL no significant difference could be discerned (Fig. 4C). However, PSAD decreased to 0.76 and 0.72 fold in NT12 and T7 respectively compared to the WT, and to 0.4 and 0.18 fold for PSAE (Fig. 4C). This decrease was corroborated by anti-PSAE immunoblots on whole cells (Supplementary Fig. S2B); similarly anti-AtpB and anti-PetB (cyt *b*_6_) supported the absence of differences in ATP synthase and cyt*b*_6_*f* levels. FD, which was not detected in the mass spectrometry data, was instead verified by immunoblot analysis and found to be largely unchanged in the mutants (Supplementary Fig. S2B).

**Figure 4.**
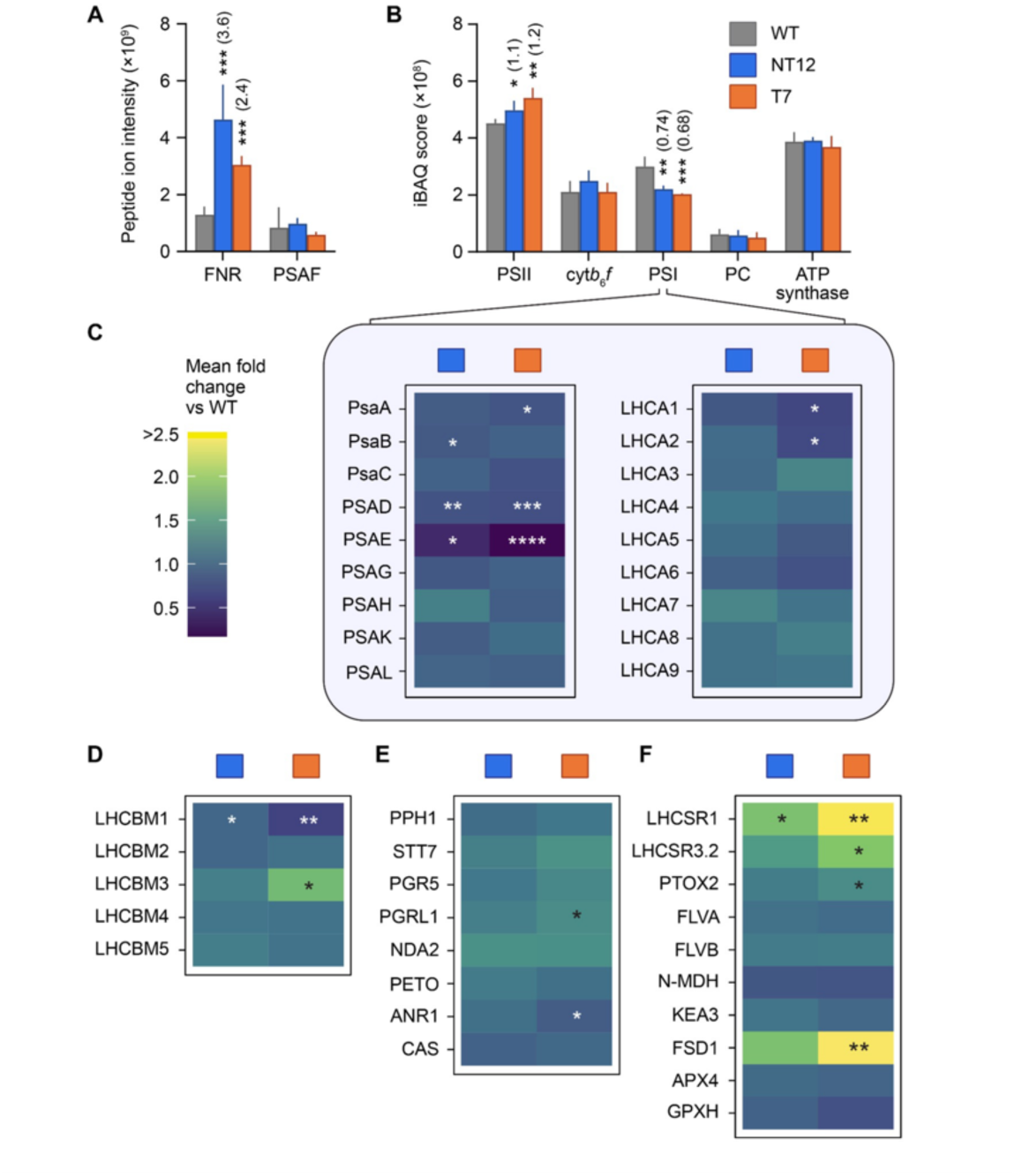
| Quantification of proteins expressed by WT, NT12 and T7 cells. A, Mass spectrometry-based label-free quantification of FNR and PSAF (sum of peptide ion intensities). **B,** Intensity based absolute quantification (iBAQ) scores for Photosystem II (mean of PsbA-D iBAQ scores, PSII), cytochrome *b*_6_*f* (mean of subunits PetA, PetB, PETC and PetD iBAQ scores, cyt*b*_6_*f*), photosystem I (mean of subunits PsaA-C, PSAD-E, PSAG-H and PSAK-L iBAQ scores, PSI) and chloroplast ATP synthase (mean of subunits AtpA, AtpB and ATPC iBAQ scores). **C,** Heat maps showing mean fold change versus WT for PSI subunits and light harvesting complex I proteins (LHCA1-9). **D,** state transition associated proteins: major light harvesting complex II proteins (LHCBM1-5). **E,** state transition associated proteins: protein phosphatase (PPH1), serine/threonine protein kinase (STT7) and cyclic electron transport (CET) associated proteins: PGR5, PGR5-like protein 1 (PGRL1), NADH-plastoquinone reductase A2 (NDA2), cyt*b*_6_*f* subunit PETO, Anaerobic Response Regulator (ANR1), calcium sensor kinase (CAS). **F,** stress associated proteins: LHCSR1, LHCSR3.2, plastid terminal oxidase 2 (PTOX2), flavodiiron proteins FLVA and FLVB, NADP-malate dehydrogenase (N-MDH), potassium efflux antiporter 3 (KEA3), superoxide dismutase FSD1, ascorbate peroxidase APX4, glutathione peroxidase GPXH. Bars show means ± SDs of quantified proteins extracted from WT, NT12 and T7 cells. Means ± SDs for proteins in panels C-F are shown in Supplementary Fig. S3. *Q* values derived from modified t-tests incorporating permutation-based false discovery rate with 250 randomizations (*n* = 4) are shown as 0.01-0.05 (*), 0.001-0.01 (**), 0.0001-0.001 (***) and < 0.0001 (****), with exact *q* values listed in Source data. Fold change relative to WT is shown in parentheses in panels A and B for differences with *q* ≤ 0.05. Differences with *q* ≥ 0.05 (not significant) are not shown.

The PSI light harvesting proteins LHCA1-9 were detected and, of these, only LHCA1 and LHCA2 were significantly different, at 0.66 fold compared to the WT in the T7 mutant (Fig. 4C). Of the PSII light harvesting proteins LHCBM1 was decreased to 0.73 and 0.61 fold compared to WT in the NT12 and T7 mutants (Fig. 4D). LHCBM2, LHCBM4 and LHCBM5 did not differ significantly in the mutants, while LHCBM3 was increased 1.7 fold in the T7 mutant but was not significantly affected in the NT12 mutant (Fig. 4D). The decrease in LHCBM1 and increase in LHCBM3 is interesting since this appears as the major phosphorylated LHCII species in the PSI-LHCI-LHCII supercomplexes (Huang et al. 2021), which are more abundant in the mutants (Fig. 3A). The state transition kinase STT7 and its cognate phosphatase were not significantly different however (Fig. 4E).

The CET-related proteins were also largely unaffected in the mutants, with no significant change in PGR5, NDA2, PETO and CAS proteins observed (Fig. 4E). Nonetheless changes were observed in the abundance of PGRL1 and ANR1 which were increased by 1.3 fold and decreased to 0.85 fold, respectively, in only the T7 mutant (Fig. 4E). Once again, the differences in PGRL1 levels were corroborated by immunoblotting of whole cells (Supplementary Fig. S2B). Changes in the LHCSR1 photoprotective protein involved in NPQ in Chlamydomonas were also detected with levels 1.8 fold and 2.5 fold higher in NT12 and T7 respectively, while LHCSR3.2 was 1.9 fold higher in T7 (Peers et al. 2009) (Fig. 4F). The plastid terminal oxidase (PTOX) protein involved in the photoprotective oxidation of PQ and transfer of electrons to water was also increased in the T7 mutant by 1.3 fold (Fig. 4F) (Nawrocki et al. 2018). In contrast, no significant differences were observed in the abundance of PCET proteins FLVA or FLVB or the NADPH-dependent malate dehydrogenase enzyme, which might be involved in reductant export from the chloroplast as in vascular plants, though direct evidence is missing in Chlamydomonas (Alric and Johnson 2017) (Fig. 4F). The putative thylakoid H^+^/K^+^ antiporter KEA3 was also unaffected in the mutants (Fig. 4F). However, levels of the superoxide dismutase (FSD1) were raised significantly in T7, by 2.7 fold, compared to the WT (Fig. 4F). No differences were observed between the mutants and WT of the ascorbate peroxidase (APX4) or glutathione peroxidase (GPXH) (Fig. 4F).

Of the CO_2_ fixation and CCM machinery, the levels of the large and small Rubisco subunits (RbcL and RBCS) were unchanged, as was the EPCY1 linker protein which binds them into the pyrenoid (Supplementary Fig. S3G). A 1.3 fold increase in the levels of the luminal carbonic anhydrase 3 (CAH3) (Mackinder 2018) responsible for converting bicarbonate into CO_2_ was observed in both mutants, while in NT12 a 1.3 fold increase in the abundance of the low carbon inducible protein LCI9 (Mackinder et al. 2017) was also seen (Supplementary Fig. S3G). The thylakoid bestrophins (BST1, BST2 and BST3) responsible for bicarbonate import across the membrane (Mukherjee et al. 2019) were all unaffected (Supplementary Fig. S3H). Among the major CBB cycle enzymes only sedoheptulose bisphosphatase levels were changed, increasing by 1.6 fold in T7 (Supplementary Fig. S3I). The major proteomic differences in the mutants therefore appear to centre around altered PSI/PSII ratio and the disruption to PSAD and PSAE levels, though differences are also seen in LHCII, photoprotection, CET and CCM related proteins.

### The chimeric PSAF-FNR mutants exhibit a CO_2_- and O_2_-dependent electron transfer phenotype

The poorer growth on minimal media compared to the WT suggests an impaired ability to fix CO_2_. This led us to investigate how photosynthesis and photoprotection were affected in the mutants. Chlorophyll fluorescence measurements were carried out with and without the addition of 10 mM sodium bicarbonate (i.e. high and low CO2) and we also tested the effect of anoxic conditions. Under high CO2/ oxic conditions, the PSII quantum yield (Y(II)) was lower in the mutant cells compared to the WT between 187 and 532 μmol photons m^-2^ s^-1^ (Fig. 5A) and PSII acceptor side limitation (1-qP) was higher at 187 and 282 (Fig. 5B). NPQ was also decreased in the mutants relative to the WT under these conditions (Figs. 5C and D). In contrast, under low CO_2_/ oxic conditions, at 187 and 282 μmol photons m^-2^ s^-1^ Y(II) was higher and 1-qP lower in T7 but similar in NT12 and WT, except at 57 μmol photons m^-2^ s^-1^ where Y(II) was higher and 1-qP lower in WT than in T7 or NT12 (Fig. 5E and F). NPQ under low CO_2_/ oxic conditions was now higher in both mutants at 532 and 702, and also higher in NT12 at 187 and 282 μmol photons m^-2^ s^-1^ than the WT (Fig. 5G and H). Y(II) was higher in T7 than NT12 or WT under high CO_2_/ anoxic conditions at 187, and 1-qP was higher in the WT at 57, 187 and 282 compared to the mutants (Figs. 5I and J). NPQ was also higher in the mutants under high CO_2_/ anoxic conditions compared to the WT (Figs. 5K and L). We tested whether the NPQ was of the ΔpH-dependent rapidly-relaxing qE variety by checking its relaxation in a subsequent dark period (Supplementary Fig. S4A). Under low CO_2_/ oxic conditions, ∼65% of NPQ was of the qE type in both mutants and WT, however in high CO_2_/ oxic conditions only ∼50% was qE in the WT and ∼20% in the mutants, consistent with a greater contribution of state transitions to the quenching (Supplementary Fig. S2A). The data suggest a CO_2_ concentration-dependent phenotype with respect to NPQ, which we confirmed in the T7 mutant, with higher concentrations of CO_2_ lowering the NPQ while showing less effect in the WT (Supplementary Fig. S4B). We therefore conclude that under stress conditions that were previously shown to favour CET, such as anoxia and low CO_2_, the NT12 and T7 mutants show an advantage in terms of NPQ generation and in line with this lower PSII acceptor side limitation compared to the WT. We confirmed this NPQ phenotype under low CO_2_ was shared by the additional independent lines we analysed (Supplementary Fig. S4C).

**Figure 5.**
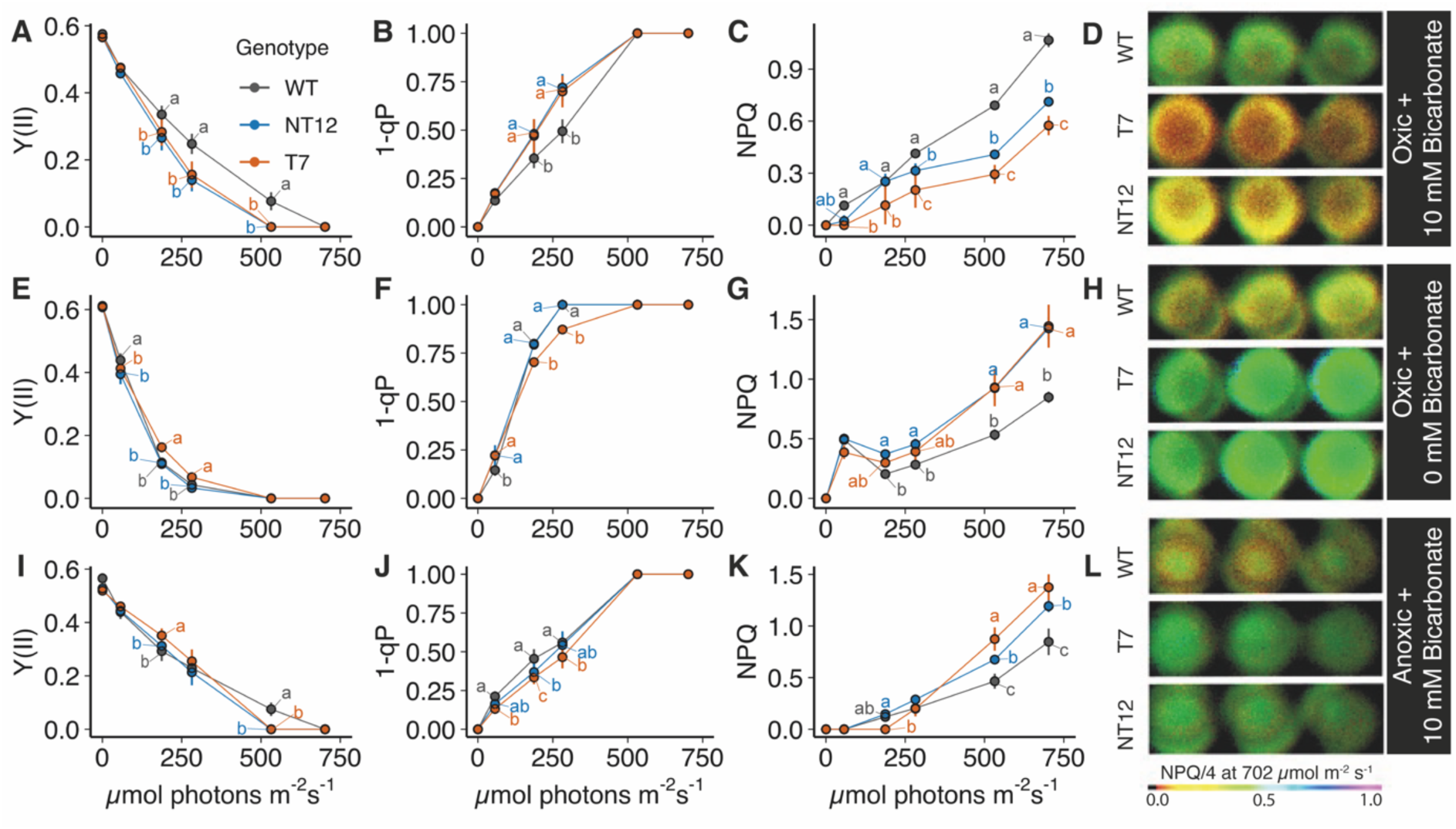
Comparison of light intensity, oxygen and CO_2_ dependence of chlorophyll fluorescence parameters of NT12 and T7 cells relative to WT. **A,E,I**, PSII quantum yield (Y(II)). **B**,**F**,**J**, Relative redox state of PSII acceptor Q_A_ (1-qP). **C,G,K**, Non-photochemical quenching (NPQ). **D,H,L**, Representative chlorophyll fluorescence image of NPQ in *Chlamydomonas* cells suspended in a well (NPQ is shown on a scale where the total value is divided by 4 to scale them between 0 and 1). In **A**-**D**, oxic cells contained 10 mM NaHCO_3_; **E**-**H,** oxic cells without addition of NaHCO_3_; **I**-**L**, anoxic cells (contained glucose oxidase, catalase and covered with clear qRT PCR film) contained 10 mM NaHCO_3_. Error bars are S.D. (at least replicates per strain). Means were compared between genotypes at each light intensity using an ordinary two-way ANOVA and corrected using Tukey’s multiple comparison test followed by a HSD test with alpha = 0.05. Different letters indicate significant differences between data points at each light intensity.

### The PSAF-FNR mutants show enhanced pmf generation and slower NADPH formation

The lower Y(II) and higher 1-qP under high CO_2_/ oxic conditions in the mutants could be explained by a limitation in the generation of ATP for CO_2_ fixation due to lower pmf, resulting in a more reduced electron transfer chain. The amplitude of the pmf, as assessed using the ECS method was somewhat lower in the WT compared to the mutants under high CO_2_/ oxic conditions at light intensities above 250 μmol photons m^-2^ s^-1^ and therefore it is unlikely that ATP is limiting for CO_2_ fixation (Fig. 6A). A larger pmf was also observed in the mutants under low CO_2_/ oxic conditions above 250 μmol photons m^-2^ s^-1^ (Fig. 6B). Under high CO_2_/ anoxic conditions, the mutants were able to generate significantly more pmf than in the WT and T7 had more pmf than NT12 at the two highest light intensities (Fig. 6C). In principle, a higher pmf could be due to either lower proton conductivity (gH^+^) as a result of downregulation of ATP synthase activity or higher proton flux across the membrane (vH^+^). The gH^+^ values were significantly higher in the WT compared to both mutants at the two highest light intensities under high CO_2_/ oxic conditions (Fig. 6D). However, under low CO_2_/ oxic and high CO_2_/ anoxic conditions the gH^+^ was more similar (Fig. 6E and F). No differences were seen for vH^+^ under high CO_2_/ oxic between mutants and WT (Fig. 6G). In contrast, higher vH^+^ was observed in both NT12 and T7 versus WT under low CO_2_/ oxic and high CO_2_/ anoxic conditions (Fig. 6H and I). The pmf and vH^+^ parameters were also significantly increased in T7 relative to NT12 under high CO_2_/ anoxic conditions at higher light (Fig. 6C and I). No major difference in either the relative amplitude or partitioning of the pmf components ΔpH and Δψ (membrane potential) were observed among the WT and mutants (Supplementary Fig. S5), consistent with unchanged KEA3 levels (Fig. 4F). Collectively, these data indicate that the mutants do not show any impairment in the ability to generate pmf and indeed have increased proton pumping (vH^+^) activity compared to the WT in high light or stress conditions.

**Figure 6.**
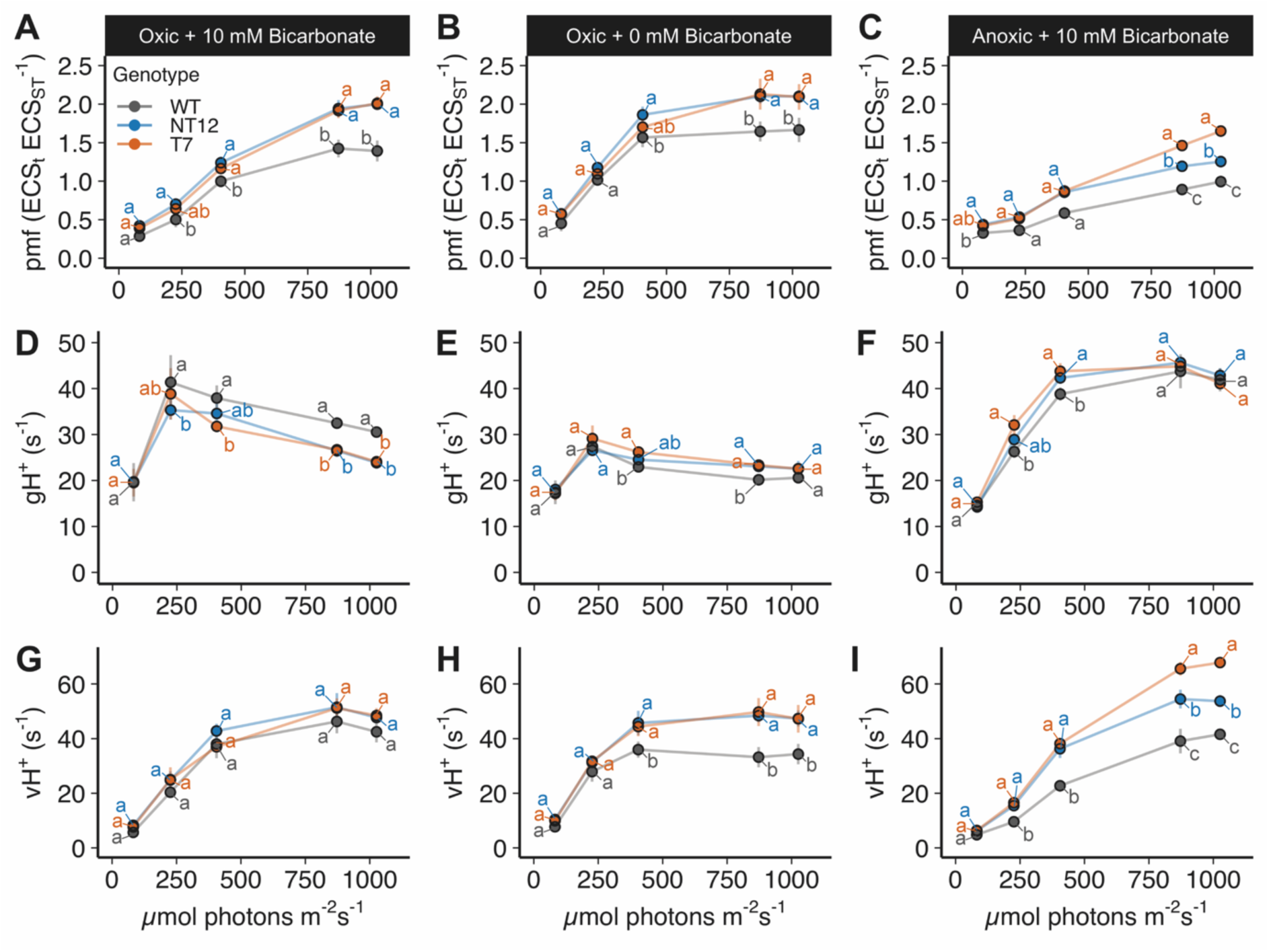
| Comparison of ECS absorption parameters of NT12 and T7 cells relative to WT. Proton motive force (pmf) under **A,** high CO_2_/ oxic**, B,** low CO_2_/ oxic and **C,** high CO_2_/ anoxic conditions. Proton conductivity (gH^+^) under **D,** high CO_2_/ oxic**, E,** low CO_2_/ oxic and **F,** high CO_2_/ anoxic conditions. Proton flux (vH^+^) under **G,** high CO_2_/ oxic**, H,** low CO_2_/ oxic and **I,** high CO_2_/ anoxic conditions. Anoxia was achieved as described in Fig. 5 except that mineral oil was used. Error bars are S.D. (at least replicates per strain). Means were compared between genotypes using an ordinary two-way ANOVA and corrected using Tukey’s multiple comparison test followed by a HSD test with alpha = 0.05. Different letters indicate significant differences between data points at each light intensity.

An alternative explanation for the slower photoautotrophic growth and lower Y(II) under high CO_2_/ oxic conditions is that the mutants have an impaired ability to reduce NADP^+^ to NADPH due to the tethering of FNR to PSI. We investigated this by monitoring the rate of NADPH fluorescence emission rise at 330 nm provoked by illumination at 500 μmol photons m^-2^ s^-1^ of whole cells (Fig. 7A). The rate of NADPH formation was ∼20% faster in each condition in the WT relative to the mutants (Fig. 7B). These data demonstrate that NADPH formation is limited in the mutants due to the slower rate of its production. In principle, slower NADPH production could prolong the lifetime of reduced FD and thus lead to enhanced ROS generation at PSI. We tested this by examining the H_2_O_2_ production. In the WT and the mutants the H_2_O_2_ production was higher under low CO_2_ compared to high CO_2_, yet the mutants produced less H_2_O_2_ than the WT under each condition (Supplementary Fig. S6).

**Figure 7.**
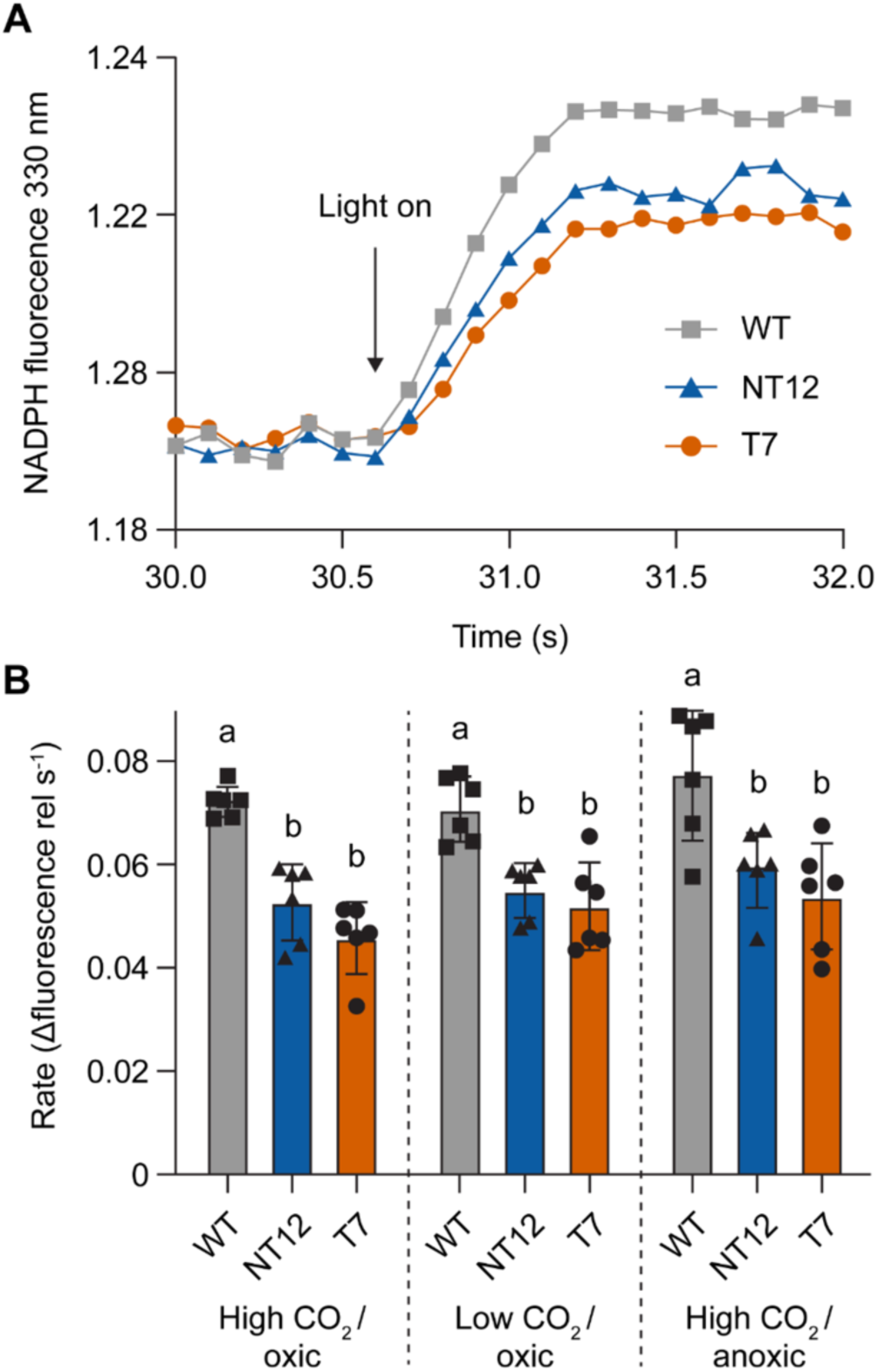
| NADPH fluorescence parameters of NT12 and T7 cells relative to WT. A, Representative traces showing response of NADPH fluorescence to actinic illumination. Light intensity was 550 μmol photons m^-2^ s^-1^. **B**, Relative rate of change in NADPH fluorescence emission intensity upon illumination. Means were compared between genotypes using an ordinary two-way ANOVA and corrected using Tukey’s multiple comparison test followed by a HSD test with alpha = 0.05. Different letters indicate significant differences between data points.

### The chimeric PSAF-FNR mutants show enhanced PSI acceptor side limitation and increased CET activity under anoxia

We next investigated the effect of the slower NADPH reduction rate on PSI activity in the mutants versus WT under high CO_2_/ oxic and high CO_2_/ anoxic conditions (Fig. 8A). In the WT under high CO_2_/ oxic conditions at 520 μmol photons m^-2^ s^-1^ around 35% of PSI RCs were donor-side limited (YND) and a further 62% could be photo-oxidised with a saturating pulse (YI), the remaining 3% was acceptor-side limited (YNA). In the NT12 mutant the YND was similar to the WT, though the YNA was significantly higher and the YI significantly lower (Fig. 8A). In the T7 mutant, YND was lower and YNA higher than the WT. Under high CO_2_/ anoxic conditions, the phenotypes of the mutants were more exaggerated with lower YND and YI and much higher YNA compared to the WT (Fig. 8A). The data are therefore consistent with the slower rate of NADPH reduction in the mutants leading to an increased acceptor side limitation of PSI.

**Figure 8.**
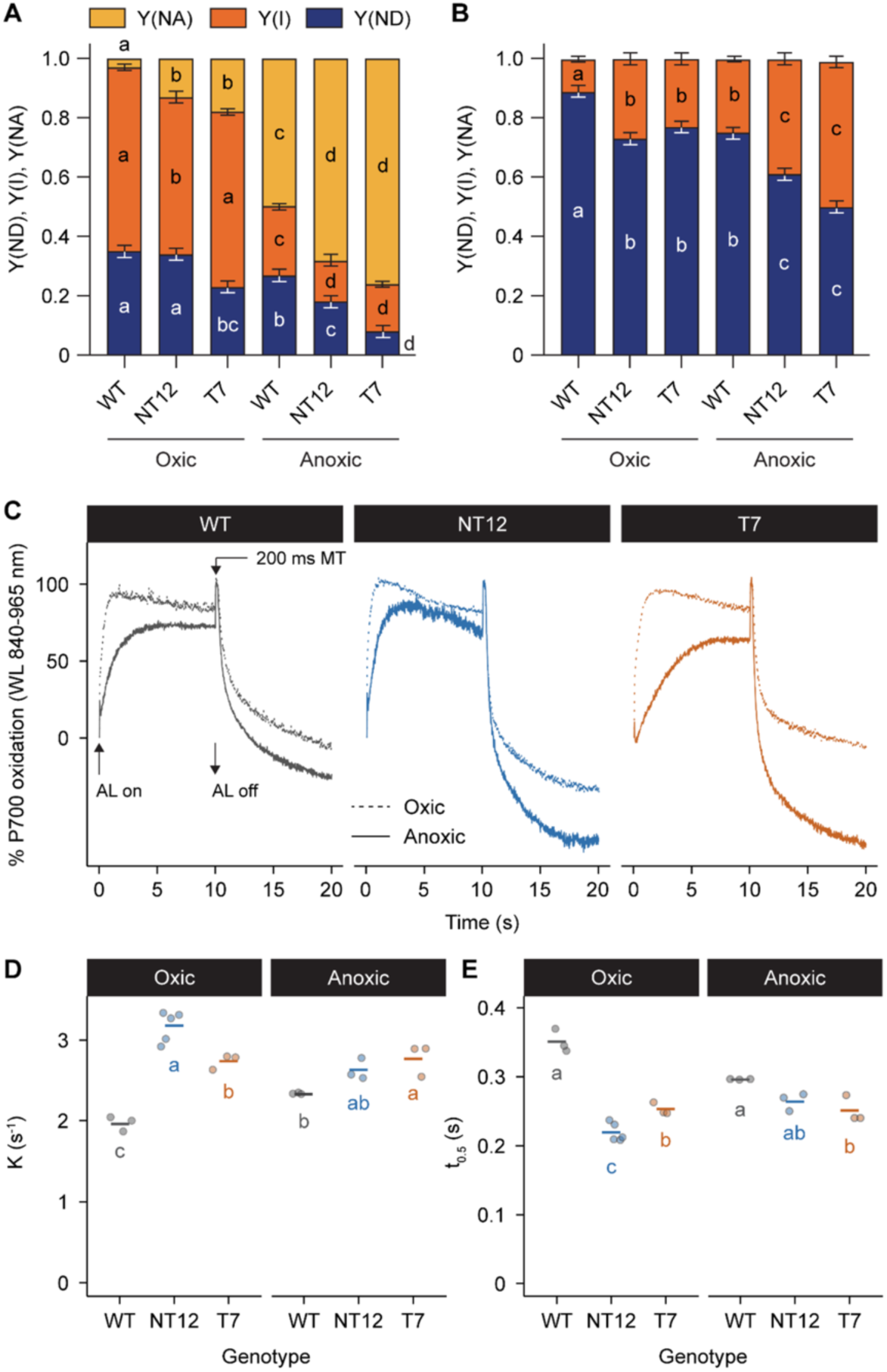
| Comparison of PSI absorption parameters of NT12 and T7 cells relative to WT. P700 redox state determined in cells subjected to 10 s illumination at 520 μmol photons m^-2^ s^-1^ red actinic light under oxic or anoxic conditions. Y(NA) = fraction of acceptor side limited PSI, Y(ND) = fraction of donor side limited PSI, Y(I) = fraction of photochemically active PSI in **A**, untreated cells and **B**, cells treated with 10 µM 3-(3,4-dichlorophenyl)-1,1-dimethylurea (DCMU) and 1 mM hydroxylamine (HA). **C**, Average raw kinetics traces of P700 oxidation/ reduction of cells treated with DCMU and HA and illuminated for 10 s with 520 µmol photons m^-2^ s^-1^ red actinic light. The calculated first order rate constant, K, and half-time, t_0.5_, of P700 reduction are shown in **D** and **E**, respectively. Horizontal lines represent means and symbols raw data points. Means were compared between genotypes using an ordinary two-way ANOVA and corrected using Tukey’s multiple comparison test followed by a HSD test with alpha = 0.05. Different letters indicate significant differences between genotypes.

We speculated that the increased pmf and vH+ in the mutants (Fig. 6A-C) might reflect an unexpectedly increased capacity for CET. We tested this further by investigating PSI properties under conditions where LET is inhibited in the presence of DCMU and HA. In the presence of these inhibitors under high CO_2_/ oxic conditions, PSI develops a strong donor side limitation (high YND) together with a small YI and no YNA (Fig. 8B). On the other hand, the mutants both showed a significant increase in YI and decrease in YND compared to the WT (Fig. 8B). When oxygen was removed (high CO_2_/ anoxic conditions), YND decreased in the WT and YI increased compared to oxic conditions (Fig. 8B). However, the two mutants showed a larger increase in YI and larger decrease in YND under anoxia (Fig. 8B). We confirmed the rate of re-reduction of PSI in the presence of DCMU and HA at 520 µmol photons m^-2^ s^-1^ was higher in the mutants compared to the WT under CET conditions (Fig. 8C-E).

We used the ECS method to exclude the possibility that increased acceptor side limitation enhanced charge recombination in the mutants, thereby accelerating PSI re-reduction under CET conditions. We assessed the ECS decay rate in the presence of DCMU and HA to inhibit LET under high CO_2_/ oxic and high CO_2_/ anoxic conditions at 520 μmol photons m^-2^ s^-1^ (Fig. 9A and B). Consistent with higher CET activity the rate of proton flux (vH^+^) was faster under anoxia in the two mutants compared to the WT, but only significantly different for T7 under oxic conditions (Fig. 9C). We confirmed that the CET phenotype was replicated in the other independent chimeric lines (Supplementary Fig. S7). Previously in Chlamydomonas, in vitro electron transfer from FD to the *b*-hemes in cyt*b*_6_*f* within the putative CET supercomplex was found to be insensitive to the inhibitor antimycin A (AA), which inhibits the PGR5 pathway in plants (Munekage et al. 2004; Iwai et al. 2010). However, recently it was shown that PGR5-dependent enhancement of *b*-heme oxidation in vivo under anoxia in Chlamydomonas is sensitive to AA (Buchert et al. 2022). If increased NPQ and pmf generation under anoxia reflect such an enhanced CET through cyt*b*_6_*f*, then logically they should be diminished by AA treatment. This was indeed the case with both WT and mutant NPQ and pmf generation significantly retarded by AA (Supplementary Fig. S8A and B). However, the differences between WT and mutants were not completely eliminated by AA. The changes in NPQ and pmf were observed without significant decreases in the overall rate of measured PSII electron transfer (Supplementary Fig. S8C) indicating AA did not retard LET at these concentrations. Collectively these data confirm that CET under anoxic conditions is in fact unexpectedly enhanced in the mutants.

**Figure 9.**
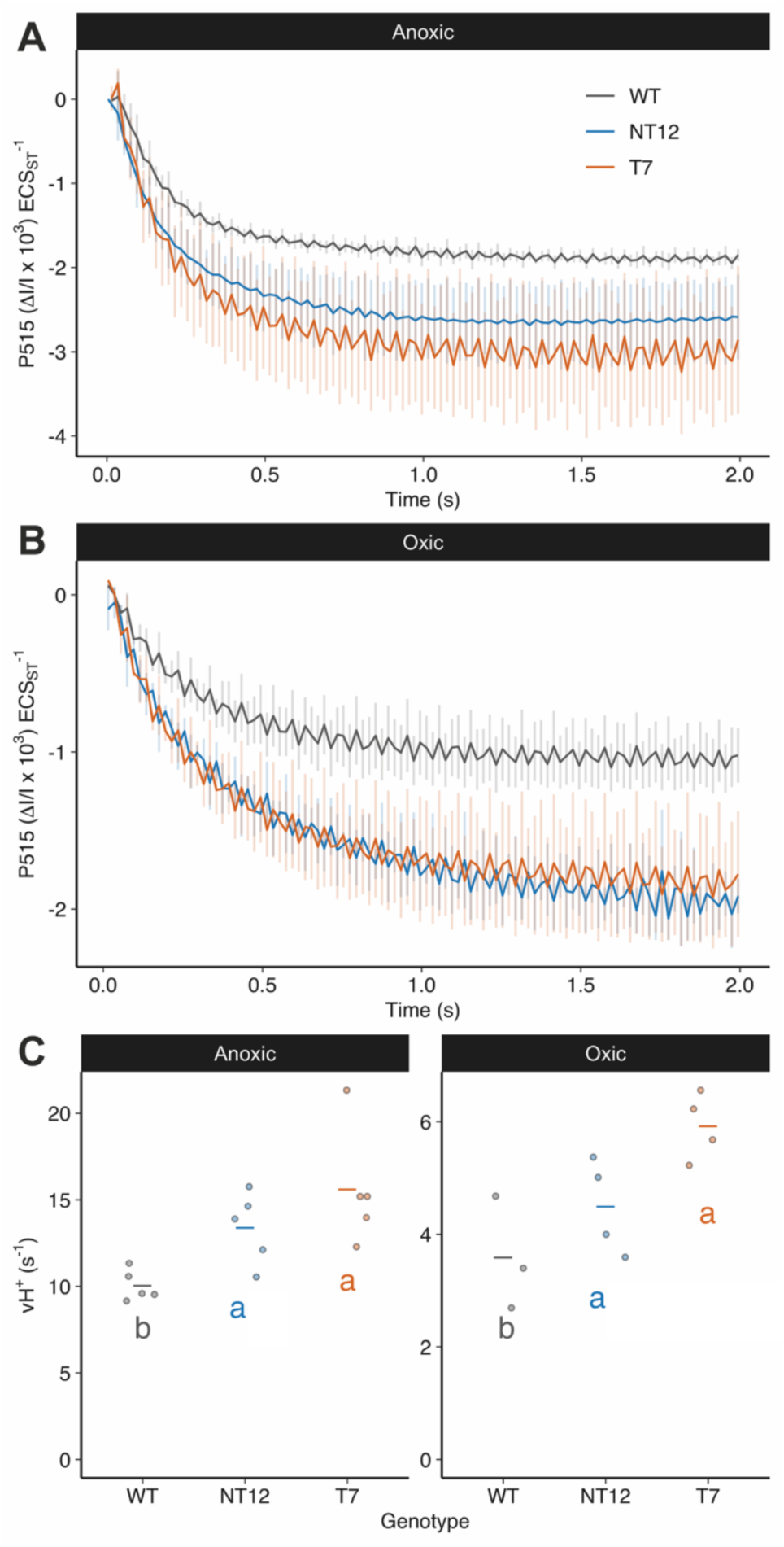
| Rate of ECS absorption decay under CET conditions in NT12 and T7 cells relative to WT. Initial rate of ECS decay in WT, NT12 and T7 cells in the presence of 10 μM DCMU and 1 mM HA under **A**, anoxic or **B**, oxic conditions upon cessation of a 10 second illumination period using 502 μmol photons m^-2^ s^-1^. **C**, Calculated initial rate (vH^+^) of ECS decay under the same conditions. Horizontal lines represent means and symbols raw data points. Means were compared between genotypes using an ordinary two-way ANOVA and corrected using Tukey’s multiple comparison test followed by a HSD test with alpha = 0.05. Different letters indicate significant differences between data genotypes.

## Discussion

In this study we tested the hypothesis that FNR location determines the allocation of photosynthetic electrons between LET and CET, with binding of FNR to PSI suggested to promote LET and binding to cyt*b*_6_*f* promoting CET (Joliot and Johnson 2011). Using CRISPR-Cas9 gene editing we successfully generated Chlamydomonas mutants where the FNR was tethered to the stromal side of PSI via PSAF. As a control we compared these tethered-only (T) mutants to both the WT and mutants which possessed both the tethered and native FNR (NT). Normally in Chlamydomonas FNR is found loosely bound to the membrane or free, with a fraction co-migrating with PSI and some with cyt*b*_6_*f* (Mosebach et al. 2017; Buchert et al. 2018). Here, the chimeric PSAF-FNR protein was found to be associated with both PSI-LHCI and the larger PSI-LHCI-LHCII supercomplex following membrane solubilisation and sucrose gradient ultracentrifugation (Fig. 3B). The chimeric protein appeared to be largely stable and did not undergo significant proteolytic cleavage in the mutants (Fig. 1F). We did however observe a perturbation in both the T and NT mutants in both overall levels of the PSI complex and specifically the stromal PSAD and PSAE subunits. As can be seen in Fig. 1A the predicted position of the chimeric PSAF-FNR at the stromal face of PSI is adjacent to PSAD and PSAE and therefore it is possible that its presence may sterically inhibit their binding. The T and NT mutants showed a lower rate of photosynthetic growth and lower PSII electron transport rate compared to the WT (Figs. 2 and 5) with increased PSII (1-qP) and PSI (YNA) acceptor side limitation, under high CO_2_ conditions (Figs. 5A-C and 8A). We traced this over-reduction of the electron transport chain in the mutants to a slower rate of NADPH formation, indeed the steady state amplitude of NADPH fluorescence was also lower (Fig. 7A).

How could the presence of the chimeric FNR disrupt NADP^+^ reduction? On the stromal/ acceptor side of PSI, the expected location of PSAF-FNR was adjacent to the PsaC bound terminal 4Fe4S clusters F_A_ and F_B_ involved in electron transfer to FD (Su et al. 2019; Naschberger et al. 2022). The reaction cycle of FNR requires the binding and unbinding of two successive FD molecules, with a reduced semiflavin storing the first electron prior to the two-electron reduction of NADP^+^ (Carrillo and Ceccarelli 2003). It is possible that the chimera delays the arrival of the second electron by ensuring each FNR is mostly only served by the immediately adjacent PSI. Interestingly, the distribution of the native FNR in the sucrose gradients was different in NT12 and the WT, with the former following the distribution of PSI (Fig. 3B). FNR is commonly found as a dimer, with dimerisation promoted in vascular plants by TIC62 and TROL, its membrane tethers (Twachtmann et al. 2012; Kramer et al. 2021). In a similar way it is possible that the tethering of FNR to PSAF promotes the dimerisation of the native FNR with it at the membrane surface in NT12, thus shifting its distribution relative to the WT (Fig. 3B). This may explain why despite the presence of additional native FNR in NT12 compared to T7 the phenotypes with regard to NADP^+^ reduction are similar (Fig. 7A). Slower NADP^+^ reduction could lead to an increased lifetime for unpaired electrons on either FD or the semiflavin of FNR, potentiating formation of superoxide, which is rapidly converted to the longer-lived ROS H_2_O_2_ by superoxide dismutase (Foyer and Hanke 2022). Indeed we observe enhanced levels of the superoxide dismutase in T7 mutant compared to the WT (Fig. 4F). However, this did not feed through to differences in H_2_O_2_ levels, which were unexpectedly lower in both low and high CO_2_ conditions in the mutants compared to the WT (Supplementary Fig. S6). As seen previously for FNR overexpressors, the overall higher levels of FNR in the NT12 and T7 mutants may protect them from any ROS build-up by pre-priming antioxidant defences (Kozuleva et al. 2016).

In contrast to the perturbation in PSII electron transfer and increased 1-qP under high CO_2_ conditions in the mutants, stress conditions with low CO_2_ or anoxia which are expected to favour alternative electron transfer mechanisms, significantly ameliorated these symptoms (Figs. 5E-I). Moreover, mixotrophic growth on acetate, which requires additional ATP relative to photoautotrophic growth (Johnson and Alric 2012; Lucker and Kramer 2013), also eliminated the growth differences compared to the WT (Fig. 2C). Consistent with this we observed higher rates of CET in the chimeric mutants by both P700 (Fig. 8C-E) and ECS (Fig. 9B and C) methods, increased proton flux (vH^+^) across the thylakoid membrane and correspondingly larger pmf (Figs. 6B, C, H and I). In line with the increases in CET and pmf, we observed higher NPQ in the mutants under stress conditions (Figs. 5G and K). The larger pmf and NPQ under anoxia in the mutants was sensitive to the inhibitor of the PGR5-dependent CET pathway inhibitor AA (Supplementary Figs. S7A and B). However the difference between the WT and mutants was not completely eliminated suggesting that either the PGR5-dependent CET pathway was only partially inhibited by AA in Chlamydomonas and/or there was enhanced electron flow through the NDA2-dependent CET pathway in the mutants. Alternatively, PCET via the flavodiiron proteins, which also use reduced FD as a substrate might contribute to the increased NPQ and pmf in the mutants compared to the WT, at least under low CO_2_ conditions. In most cases the NT12 and T7 lines behaved in a similar way with regard to pmf, NPQ and CET, suggesting the FNR phenotype of the chimera was dominant. Though where differences occurred they were normally more extreme in the tethered-only T7 line compared to NT12, suggesting the presence of the native FNR can partially rescue the phenotype.

Although CET and state transition can operate independently (Terashima et al. 2012; Takahashi et al. 2013), it is possible that the higher CET capacity reflects the increased shift to State II in the mutants which partitions more excitation towards PSI (Cardol et al. 2009). State II is induced by the activation of the kinase STT7 via binding of PQH_2_ to the oxidising site on the cyt*b*_6_*f* complex (Zito et al. 1999; Depège et al. 2003). The more reduced PQ pool (inferred by 1-qP, Fig. 5B) in the mutants observed in light curves under high CO_2_/ oxic conditions is consistent with activation of State II under these conditions (Fig. 3E). However, consistent with recent reports (Buchert et al. 2018), we found cyt*b*_6_*f* and PSI-LHCII distribution were poorly correlated (Fig. 3B), suggesting the association between these complexes is extremely labile or it is unnecessary for CET. Alternatively, the acceleration of CET under these conditions in the mutants could be due to pleiotropic effects of the mutation on CET effector proteins. PGRL1 levels were slightly increased in T7, though not significantly in NT12. In contrast, ANR1, a putative auxiliary subunit of cyt*b*_6_*f* involved in CET, abundance was lower than in the WT (Takahashi et al. 2016) (Fig. 4E). A decrease was also observed in LHCA2 proteins, the absence of which was previously shown to promote CET (Terashima et al. 2012; Steinbeck et al. 2018) (Fig. 4C). As stated above we found PSAD and PSAE abundance is lower in the chimeric mutants. Interestingly, past work in Arabidopsis mutants lacking PSAD and PSAE shows that while PSI remains functional the balance between LET and CET is shifted towards the latter (Hald et al. 2008b). Indeed, in both cases CET was enhanced by around 1.5 fold compared to the WT, similar to the enhancement seen here. It is possible that loss of these subunits somehow provides easier access for electrons to the CET pathway. Given the recent reports in Arabidopsis that membrane tethered FNR promotes CET (Kramer et al. 2021; Rodriguez-Heredia et al. 2021), we suggest that the FNR chimera likewise results in more electrons being channelled into the CET (and possibly PCET) pathway at the expense of LET as summarised in Figure 10. Such an explanation could rationalise why direct binding of FNR to cyt*b*_6_*f* (prevented by the chimera) is unnecessary for CET. Indeed, the decreased CET in the PGR5 mutant (Johnson et al. 2014; Buchert et al. 2020) goes along with a depletion of membrane bound FNR (Mosebach et al. 2017). Finally, it is also possible that chimera affects the conformation of FNR in such a way to enhance the relative rate of the back reaction (i.e. NADPH to FD) relative to NADP^+^ reduction. These ideas now require further testing.

**Figure 10.**
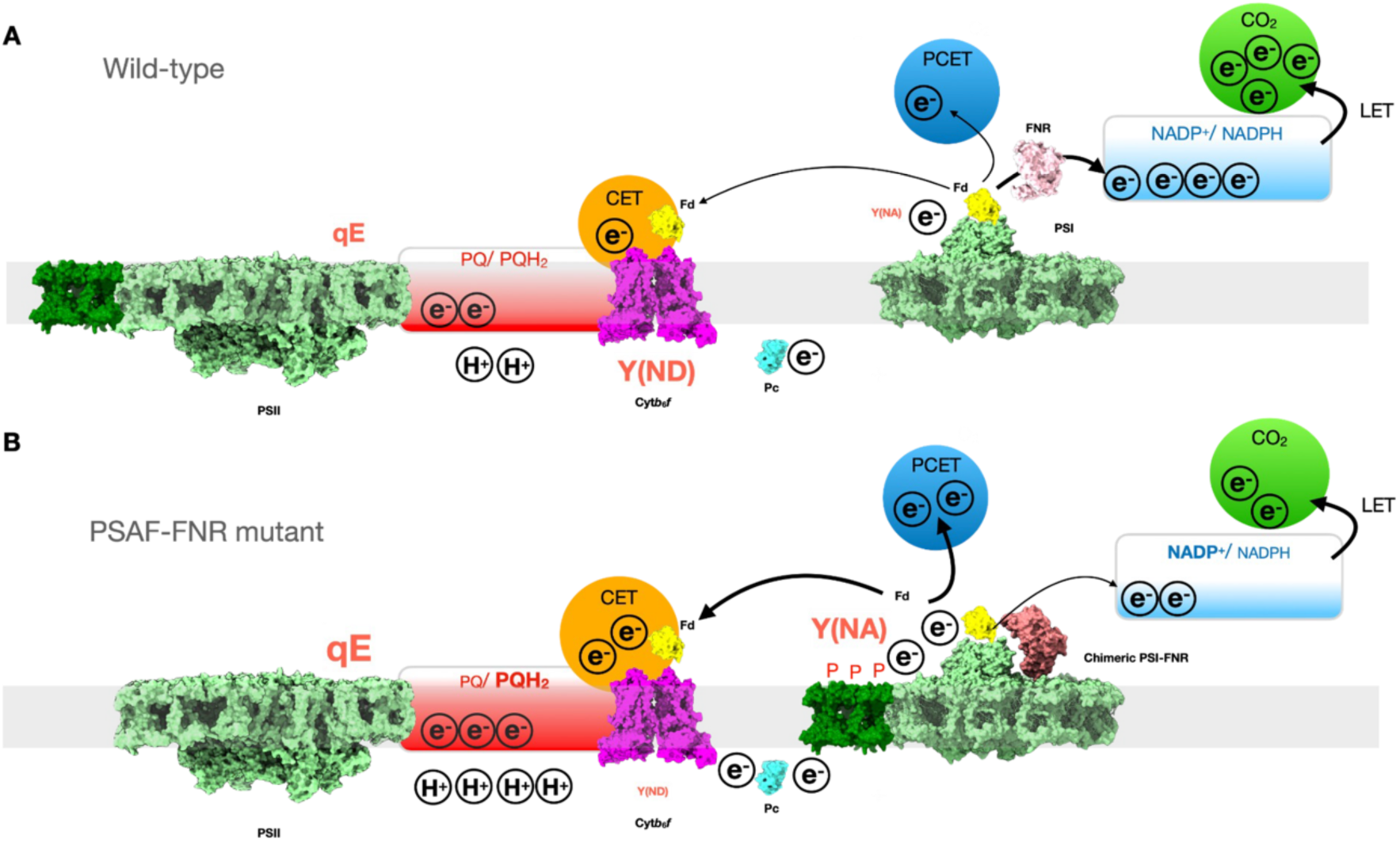
| Schematic model outlining the effect of PSAF-FNR chimera on photosynthesis. A, In the WT the native FNR efficiently shuttles electrons from FD to NADP^+^ to drive LET and CO_2_ fixation. Alternative electron sinks such as CET and PCET receive sufficient electrons from FD to support additional ATP synthesis for CO_2_ fixation. The ΔpH and redox poise of the NADP^+^/ NADPH pool support a moderate level of qE and allow oxidation of PSI (Y(ND)). PSI acceptor side limitation (Y(NA)) is minimal. **B**, Contrastingly in the PSAF-FNR mutant the tethering of FNR to PSI perturbs the forward ET reaction from FD to NADP^+^ leading to accumulation of reduced FD, increased PSI acceptor side limitation (Y(NA)) and increased PQ pool reduction, driving a transition to state II. Increased availability of reduced FD stimulates CET and likely PCET, increasing ΔpH which increases qE. However, Y(ND) is not stimulated in the mutant, possibly due to loss of NADP^+^/NADPH redox poise.

Despite higher pmf and NPQ, we observed lower P700 oxidation (YND) under anoxia in the chimeric mutants compared to the WT (Fig. 8). Since the fraction of pmf present as ΔpH was essentially unchanged in the mutants (Supplementary Fig. S5), consistent with unchanged KEA3 levels (Fig. 4F), this suggests that another factor is at play in suppressing YND. Saliently, a similar phenotype was recently reported for the Arabidopsis *hope2* mutant, which showed increased CET activity and normal NPQ, though higher YNA and lower YND than the WT (Degen et al. 2023). Increasing the flow of electrons to the PSI donor side via high CET may therefore mitigate the extent of any increase in YND. Alternatively, YND may require a proper poising of the NADPH pool as previously suggested (Hald et al. 2008a). Since there was no evidence for enhanced YND in the mutants it can be ruled out that the chimera affected interaction between PC and PSI at the lumenal face of PSAF (Fig. 8A).

In summary, using a CRISPR-Cas9 gene editing approach we were able to modify the location of FNR through construction of a chimera with PSAF and deletion of the native copies. This allowed us to test the hypothesis that FNR binding to PSI promotes LET over CET. Our results are inconsistent with this simple hypothesis and instead support a model where FNR location (membrane bound versus loosely bound/ free) plays a role in determining the partition of electrons between the LET and CET pathways.

## Materials and methods

### Strains and culturing conditions

The wild type Chlamydomonas strain for all experiments was UVM4, kindly provided by Prof. Ralph Bock. UVM4 was generated by Neupert and colleagues in 2009 (Neupert et al. 2009) and derives from strain CC-4350, available from the Chlamydomonas Resource Centre (https://chlamycollection.org). Cells were cultured in Tris-phosphate (TP) or Tris-acetate-phosphate (TAP) liquid media with revised trace elements (Kropat et al. 2011), pH 7.4, in flat bottomed flasks with 120 rpm of shaking in air at 20 °C under a 14 h/ 10 h day-night cycle at 60 µmol photons m^-2^ s^-1^ white LED light. All Chlamydomonas strains were maintained under low (10-25 µmol photons m^-2^ s^-1^) white fluorescent light on TAP + 1.5% agar plates containing 20 μg mL^-1^ paromomycin and/ or 25 μg mL^-1^ hygromycin where appropriate.

### Cassette construction

The *PSAF-FNR* expression cassette was assembled into a backbone from the pLM005 vector family (Mackinder et al. 2017). The ∼1.2 kbp native *PSAF* nucleotide sequence (Cre09.g412100, Phytozome v5.6) was first amplified from genomic DNA (gDNA) and inserted into pLM005 between the *PSAD* (Cre05.g238332, Phytozome v5.6) promoter + 5’UTR and terminator, downstream of the *AphVIII* cassette conferring paromomycin resistance. Next, the native *FNR* nucleotide sequence (Cre11.g476750, Phytozome v5.6) was analysed for the presence of a chloroplast transit peptide using ChloroP (Emanuelsson et al. 1999), now renamed TargetP (Armenteros et al. 2019). A ∼2.2 kbp sequence excluding the transit peptide sequence was amplified from gDNA using primers that included a 5’ linker sequence and 3’ Strep-tag II sequence, then inserted into the vector in frame with the *PSAF* sequence by restriction-ligation cloning. Lastly, a series of silent mutations were made by overlap extension PCR to remove Cas9 recognition sites in *PSAF* exon 3 and *FNR* exon 6 of the *PSAF-FNR* expression cassette. Suitable Cas9 sites were identified by searching the *C. reinhardtii* genome v5.5 using CRISPR-P 2.0 (Liu et al. 2017) and CRISPRdirect (Naito et al. 2015) and chosen due to minimal predicted off-target cleavage sites (Supplementary Table S1). Three silent mutations were made to the *PSAF* exon 3 recognition site, including removal of the PAM sequence. Codon restrictions around the *FNR* exon 6 recognition site meant the PAM sequence could not be modified, so four silent mutations were made. The final plasmid, pTE051, was sequenced to confirm correct assembly, then prepared for transformation by digestion with the blunt double-cutter Eco32I to produce the ∼7 kbp *PSAF-FNR* expression cassette, shown in Fig. 1A. Sequence information is provided in Supplementary Table S1.

### CRISPR-mediated mutation

CRISPR-mediated mutations were made to the Chlamydomonas nuclear genome using the Alt-R CRISPR system (Integrated DNA Technologies). The major route for DNA repair in Chlamydomonas is the non-homologous end-joining pathway (NHEJ), which can incorporate fragments of exogenous DNA into the breakage site during the repair process (Ghribi et al. 2020). Accordingly, to insert the *PSAF-FNR* expression cassette into the native *PSAF* gene, linearised DNA encoding the *PSAF-FNR* expression cassette (which also includes the paromomycin resistance cassette, *AphVIII*) was co-transformed into WT (UVM4) cells by electroporation with ribonucleoprotein (RNP) consisting of a modified Cas9 enzyme from *Saccharomyces pyogenes* (SpCas9) in complex with a custom single-guide RNA (sgRNA) with homology to *PSAF* exon 3. Similarly, to delete FNR, linearised DNA encoding a hygromycin resistance cassette (*AphVII*) from pLM006 (Mackinder et al. 2017) was co-transformed into NT12 cells with RNP containing sgRNA with homology to *FNR* exon 6. sgRNA sequences were designed as mentioned above, so that only the native copies of *PSAF* or *FNR* were targeted for CRISPR-mediated cleavage, while the edited *PSAF* and *FNR* sequences in the *PSAF-FNR* expression cassette were ignored by the RNP.

Briefly, cells were grown in TAP to mid-log phase at 20 °C then incubated at 33 °C for 3 h followed by harvesting and resuspension in 120 µl MAX Efficiency Transformation Reagent for Algae (Thermo Fisher) to a concentration of 2 x 10^8^ cells mL^-1^ in a 4 mm gap cuvette. Cuvettes were incubated in a water bath at 16 °C for 5 min prior to mixing of 500 ng linearised DNA and 61 pmol RNP and a single pulse at 0.8 kV in a MicroPulser Electoroporator (BioRad). Following transformation, cells were recovered at 33° overnight in TAP + 40 mM sucrose under low light (10-25 µmol photons m^-2^ s^-1^) and 120 rpm shaking before plating. Colonies were screened by colony PCR to identify disruptions at the target locus. Primers oTE587 and 588 were used to amplify across the *PSAF* gDNA coding sequence (Fig. 1B (ii)). Primers oTE497 and 511 were used to amplify a ∼1 kbp fragment from within the *PSAF-FNR* expression cassette (Fig. 1B (iii)). Primers oTE347, 348 and 024 were used in a three-primer PCR to amplify across one of the ends of the insertion site of the *AphVII* cassette in the native FNR exon 6 sequence (Fig. 1E (iii)). Primer sequences and sgRNA details are provided in Supplementary Table S1.

### Thylakoid isolation and protein purification

To prepare thylakoid membranes for sucrose gradient separation, WT and mutant cell cultures were grown in 1 L TAP medium under a 14 h/ 10 h day-night cycle at 60 µmol photons m^-2^ s^-1^ white LED light at 20 °C until the cells reached mid-log growth phase (2-4 x 10^6^ cells mL^-1^), the cells were then transferred to TP medium for 2-3 days to switch to autotrophic growth, at which point the cells were collected by centrifugation at 4000 x *g* for 5 mins. Cell pellets were washed once and resuspended in Buffer A (10 mM HEPES pH 7.5, 0.33 M sucrose, 10 mM EDTA, 10 mM NaF, 1.5 mM KCl). EDTA-free protease inhibitor tablets (Merck) and DNAse were added prior to lysis by two passes through a French press at 8000 psi. The cell lysate was centrifuged at 3000 x *g* for 10 min, 4 °C and thylakoids collected by centrifugation at 56000 x *g*, 30 min, 4 °C. Thylakoids were washed once in Buffer B (10 mM HEPES pH 7.5, 10 mM EDTA, 1.5 mM KCl) and resuspended to a chlorophyll concentration of 1 mg mL^-1^. Lauryl maltose neopentyl glycol (LMNG) was added to the thylakoids to a final concentration of 2% (w/v) and incubated for 30 min, 4 °C in the dark with gentle agitation. Insoluble material was pelleted at 56000 x *g*, 30 mins, 4 °C. The chlorophyll concentration across all samples was normalised and 500 µL of each loaded onto sucrose density gradients made by the freeze-thawing 10 mM HEPES pH 7.5, 0.85 M sucrose, 0.006% LMNG (w/v). The samples were centrifuged for 22 hours at 288000 x *g*, 4 °C. 1 mL fractions were collected by top-down unloading using a syringe and needle.

### Electrophoresis and immunoblotting

Cell quantities were normalised to chlorophyll concentration for whole cell blots and by volume for sucrose gradient fraction blots. Proteins were extracted by boiling and sonicating for 10 minutes in Laemmli buffer prior to separation by SDS-PAGE and immunoblotting. SDS-PAGE and immunoblotting were carried out as previously described (Wood et al. 2018). Antibodies were obtained from Agrisera (AtpB-AS05 085; FD-AS06 121; FNR-AS15 2909; LHCBM5-AS09 408; PetA-AS06 119; PetB-AS03 034; PetC-AS08 330; PGRL1-AS19 4311; PsaE-AS22 4731) with the exception of the PsaF antibody which is described elsewhere (Hippler et al. 1997).

### Low-temperature fluorescence emission spectroscopy (77 K)

WT, NT12 or T7 cell cultures were diluted in TP to ∼5 mg Chl mL^-1^ then frozen in liquid nitrogen and analysed using a FluoroLog FL3-22 spectrofluorimeter (Jobin Yvon). Fluorescence emission spectra were recorded by excitation at 435 nm and monitoring between 600-800 nm. Spectra were normalised to the peak at 680 nm, associated with fluorescence from PSII.

### Physiological measurements

For oxic conditions, cells were equalised in TP (100 µg Chl mL^-1^) containing 15% Ficoll, then incubated in the dark for 10 min. For anoxic conditions, cells were mixed with oxygen scavengers: 50 mM glucose, 30 U mL^-1^ catalase and 10 U mL^-1^ glucose oxidase. For high CO_2_ 10 mM sodium bicarbonate was added prior to experiments, for low CO_2_ no additions were made. Cells were then overlaid with 500 μl mineral oil or sealed with clear qRT PCR film to prevent gas exchange, and cells were incubated in the dark for 40 min without shaking to deplete oxygen.

For chlorophyll fluorescence, electrochromic shift (ECS) and P700 measurements, cells were grown in TAP medium until mid-log phase under a 14 h/ 10 h day-night cycle at 60 µmol photons m^-2^ s^-1^, white LEDs, 120 rpm shaking at 20-22 °C. Cells were then collected by centrifugation at 2100 x *g* for 2 min, resuspended in TP medium and returned to the above conditions for at least 24 h. For high light conditions, cells were placed under 200 µmol photons m^-2^ s^-1^ of red actinic light for 24 h using a magnetic stirrer and sodium bicarbonate was added to a final concentration of 10 mM. Prior to measurements, cells were collected via centrifugation and resuspended in 15% Ficoll (w/v) in TP.

Chlorophyll fluorescence measurements were performed using an ImagingPAM (Heinz Walz GmbH, Effeltrich, Germany). 400 µL of cells (100 µg Chl mL^-1^) resuspended in 15% Ficoll were added to each well of a 24-well plate (see figure legends for details of treatments). Cells were dark-adapted for 10 min prior to Fm determination using an 800 ms saturating pulse. Light curves and induction curves were performed according to the description in the figures.

Electrochromic shift measurements were performed using the DualPAM (Heinz Walz GmbH, Effeltrich, Germany) with the P515 emitter/detector module. 1 mL of cells (33 µg Chl mL^-1^) in 15% Ficoll were added to a quartz cuvette, which was placed in the sample holder so that the emitter head could be lowered into the cuvette. Cells were dark-adapted for 5 min prior to measurements for oxic conditions and 35 min for anoxic conditions, as detailed in the figure legends. Proton motive force was calculated from the decay of the P515 signal when red actinic light was turned off by fitting a single exponential decay to the first 300 ms in the dark to determine the span of the signal decay (ECS_t_). The proton conductance (gH^+^) was calculated as the inverse of the rate constant of this decay. Proton flux (vH^+^) was calculated as ECS_t_ x gH^+^. P700 oxidation was measured in a similar way using a DualKLAS (Heinz Walz GmbH, Effeltrich, Germany).

Decay of P700 oxidation (WL 840-965 nm) was measured after 10s of red actinic light illumination at 502 µmol photons m^-2^ s^-1^. At the end of the illumination period, a saturating 200 ms multiple turnover flash was used to determine maximum P700 oxidation (Pm) and the first 1.5 seconds were used to fit a nonlinear decay function for calculation of K and t_0.5_. Cells (100 µg Chl mL^-1^) were dark adapted prior to measurements with 10 µM DCMU + 1 mM HA.

### NADPH fluorescence

NADPH fluorescence was measured at 460 nm with 10 nm slit width (excitation 340 nm, 5 nm slit width), integration time 0.01s. Actinic illumination was provided by a Schott KL 1500 HAL light source with 610 nm long pass filter. Samples were illuminated for 30 s followed by 60 s of dark. Rates were calculated using a linear fit over the time in which the light was turned on or off, or during the illumination period. 15 samples were taken for each condition from 3 biological replicates.

### H_2_O_2_ assay

For H_2_O_2_ quantification, cells were grown as described above and 1 mL of TP culture was then diluted at a 1:1 ratio with fresh TP medium, containing 1 U of horseradish peroxidase and 5 μM of Amplex Red (ThermoFisher) and incubated for 1 h under growth conditions. Cells were removed by centrifugation and the fluorescence intensity of oxidised Amplex Red (resorufin) was immediately quantified using a BMG Labtech Plate Reader Excitation/Emission 560/580-10 nm and compared against a linear H_2_O_2_ standard.

### Mass spectrometry

Cells (WT, NT12 and T7 strains) were grown in TAP liquid medium as four independent replicate cultures. At mid-log phase (1-5 x 10^6^ cells mL^-1^) the cells were transferred to TP medium for 2-3 days to switch to autotrophic growth. Cells were harvested by centrifugation in 5 mL aliquots at 1200 x *g* for 2 min and the pellets flash frozen in liquid N_2_ before storage at −80 °C. Cell pellets (one per analysis) were suspended in 20 µl 2% (w/v) Na dodecylsulphate, 40 mM Tris base, 60 mM dithiothreitol (DTT) (lysis buffer) and solubilised by bead beating with an equal volume of 0.5 mm silica-zirconia beads at 95 °C (Krynická et al. 2019). The cell lysates were recovered after brief centrifugation and the beads washed with 50 µl lysis buffer which was then combined with the primary lysate. Proteins were extracted from the lysates by precipitation using a 2D clean-up kit (Cytiva) according to the manufacturer’s protocol and the pellets solubilised in 90 µl 4% (w/v) Na deoxycholate, 100 mM Tris-HCl, pH 8.5 (Duan et al. 2024). After reduction (10 mM DTT, 56 °C, 5 min) and S-methylthiolation (20 mM S-methyl methanethiosulfonate, room temperature, 10 min), proteins were digested with 1 µg premixed trypsin/endoproteinase Lys-C (Promega) at 37 °C for 16 h with shaking at 800 rpm. Peptides were isolated after precipitating the deoxycholate by adding absolute acetonitrile to 20% (v/v) and 10% (v/v) trifluoroacetic acid (TFA) to 0.5% (v/v). After dilution to 5% (v/v) acetonitrile with 3 vol 0.5% (v/v) TFA, peptides were desalted using C18 spin columns (Thermo Fisher Scientific) according to the manufacturer’s instructions and dried in a vacuum concentrator.

Quantitative proteomic analysis by nano-flow liquid chromatography coupled to mass spectrometry (nanoLC-MS) was performed using the Vanquish Neo/Exploris 480 system (Thermo Fisher Scientific) with acquisition parameters as previously described (Flannery et al. 2021) except that a 2-h gradient and 20 dependent scans were selected. Protein identification and label-free quantification (LFQ) were performed by searching the MS data files against the *Chlamydomonas reinhardtii* reference proteome database (https://www.uniprot.org/proteomes/UP000006906, downloaded on 3 October 2023 and edited to include the FNR-PsaF chimeric protein sequence) using MaxQuant v. 1.6.10.43 (Cox and Mann 2008). For LFQ, the iBAQ (Schwanhäusser et al. 2011) and ‘match between runs’ options were selected and the output was configured for export into Microsoft Excel and GraphPad Prism v. 10.2.2 using Perseus v. 2.0.10.0 (Tyanova et al. 2016).

## Data availability statement

The mass spectrometry proteomics data have been deposited to the ProteomeXchange Consortium via the PRIDE partner repository (http://proteomecentral.proteomexchange.org) with the data set identifier PXD052753. All other data can be obtained from the corresponding author upon request.

The author responsible for distribution of materials integral to the findings presented in this article in accordance with the policy described in the Instructions for Authors (www.plantcell.org) is: Matthew P. Johnson.

## Acknowledgments

M.P.J. acknowledges funding from the Leverhulme Trust grants RPG-2019-045 and RPG-2021-345. M.P.J. also acknowledges financial support from the Biotechnology and Biological Sciences Research Council (BBSRC UK), award number BB/V006630/1. T.Z.E.M, was supported by a BBSRC White Rose DTP studentship in Mechanistic Biology. F.R.H was supported by a Diamond PhD studentship. C.N.H. is supported by a European Research Council Synergy award 854126. The authors wish to thank Professor Ralph Bock for the gift of the UVM4 strain and Dr Trong Khoa Pham, biOMICS Facility of the Faculty of Science Mass Spectrometry Centre at the University of Sheffield, for conducting the nanoLC-MS and MaxQuant analyses. M.H. was supported by the DFG (Deutsche Forschungsgemeinschaft) [grant number HI 739/13-3] and DFG FOR 5573—GoPMF [grant number HI 739/25-1].

## Supplementary Data

The following materials are available in the online version of this article.

Supplementary Figure S1. 77K fluorescence emission spectra of WT, NT12 and T7 cells.

Supplementary Figure S2. ECS quantification of PSI/PSII ratio and immunoblots of key proteins.

Supplementary Figure S3. Additional quantification of proteins expressed by WT, NT12 and T7 cells.

Supplementary Figure S4. Effect of added sodium bicarbonate on NPQ in oxic cells.

Supplementary Figure S5. Partitioning of proton motive force into ΔpH and Δψ.

Supplementary Figure S6. Hydrogen peroxide levels in NT12, T7 and WT cells.

Supplementary Figure S7. Rate of ECS absorption decay under CET conditions in T and NT mutants relative to WT.

Supplementary Figure S8. Effect of Antimycin A on chlorophyll fluorescence and ECS parameters in WT, NT12 and T7 cells under anoxia with 10 mM NaHCO_3_.

Supplementary Table S1. Plasmid, sgRNA and primer sequence information.

## Author Contributions

T.E.M, M.S.P and G.E.D performed the majority of the research and analysed the data. P.J.J, K.J.R, F.R.H and F.B performed additional experiments. T.E.M, M.S.P, G.E.D, C.N.H, A.H, M.H. and M.P.J conceived the experiments and wrote the manuscript. All authors read and approved the manuscript prior to submission. T.E.M, M.S.P and G.E.D contributed equally to the manuscript. All authors agreed that they can list the paper as Emrich-Mills et al., Proctor et al. or Degen et al. for the purpose of their CVs.

