## Supplemental Data Figures 1-8 for "Tethering ferredoxin-NADP^+^ reductase to photosystem I promotes photosynthetic cyclic electron transfer"

### Supplemental Fig 1

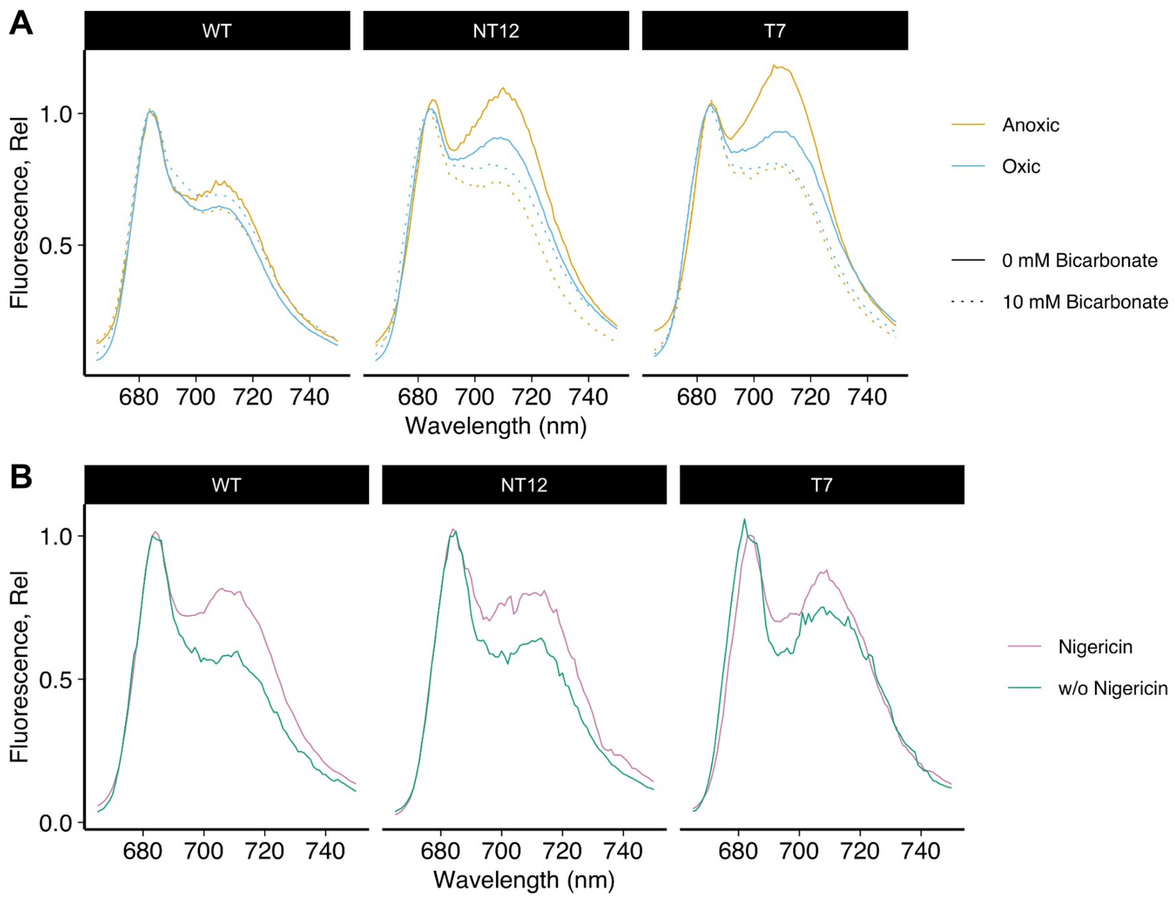


**Supplementary Figure S1 | 77K fluorescence emission spectra of WT, NT12 and T7 cells illuminated with 420 nm light. A,** Four conditions are shown, with high/ low $CO_2$ indicating 0 or 10 mM added sodium bicarbonate and oxic/ anoxic indicating with/ without 50 mM glucose, 30 U mL$^{-1}$ catalase and 10 U ml$^{-1}$ glucose oxidase. **B,** Effect of inclusion or omission of 10 µM nigericin on high $CO_2$/ anoxic cells. Supports Fig. 3.

### Supplemental Fig 2

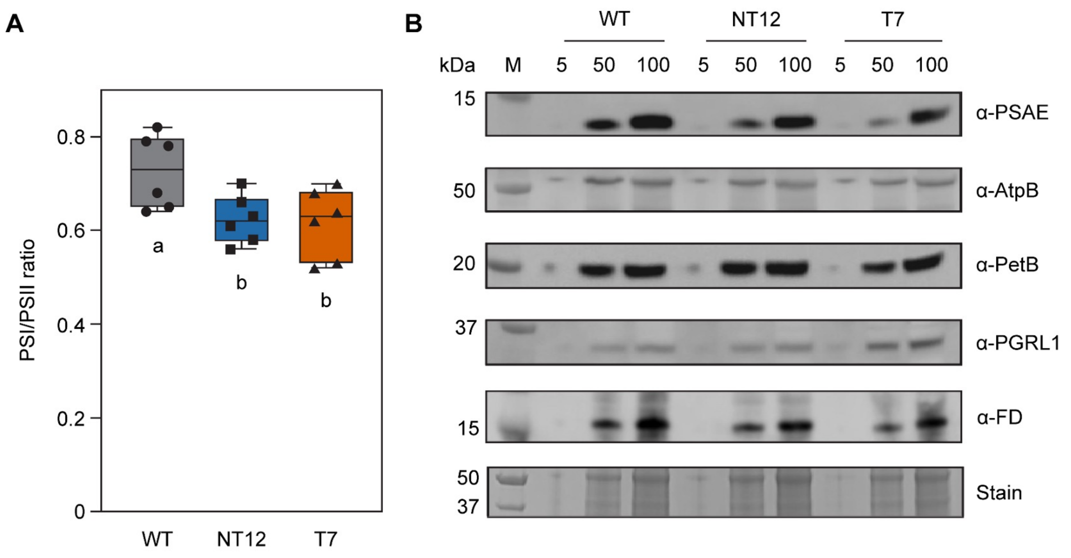


**Supplementary Figure S2 | A, ECS quantification of PSI/PSII ratio. B, Immunoblots of key proteins.** Blots were performed on whole cells of WT, NT12 and T7 using antibodies to PSI (PSAE), ATP synthase β subunit (AtpB), cytochrome $b_6$ (PetB), PGRL1 and FD. Coomassie Stain shows equal protein loading at 5, 50 and 100 μg Chl $mL^{-1}$. M is the molecular weight marker. Supports Fig. 4.

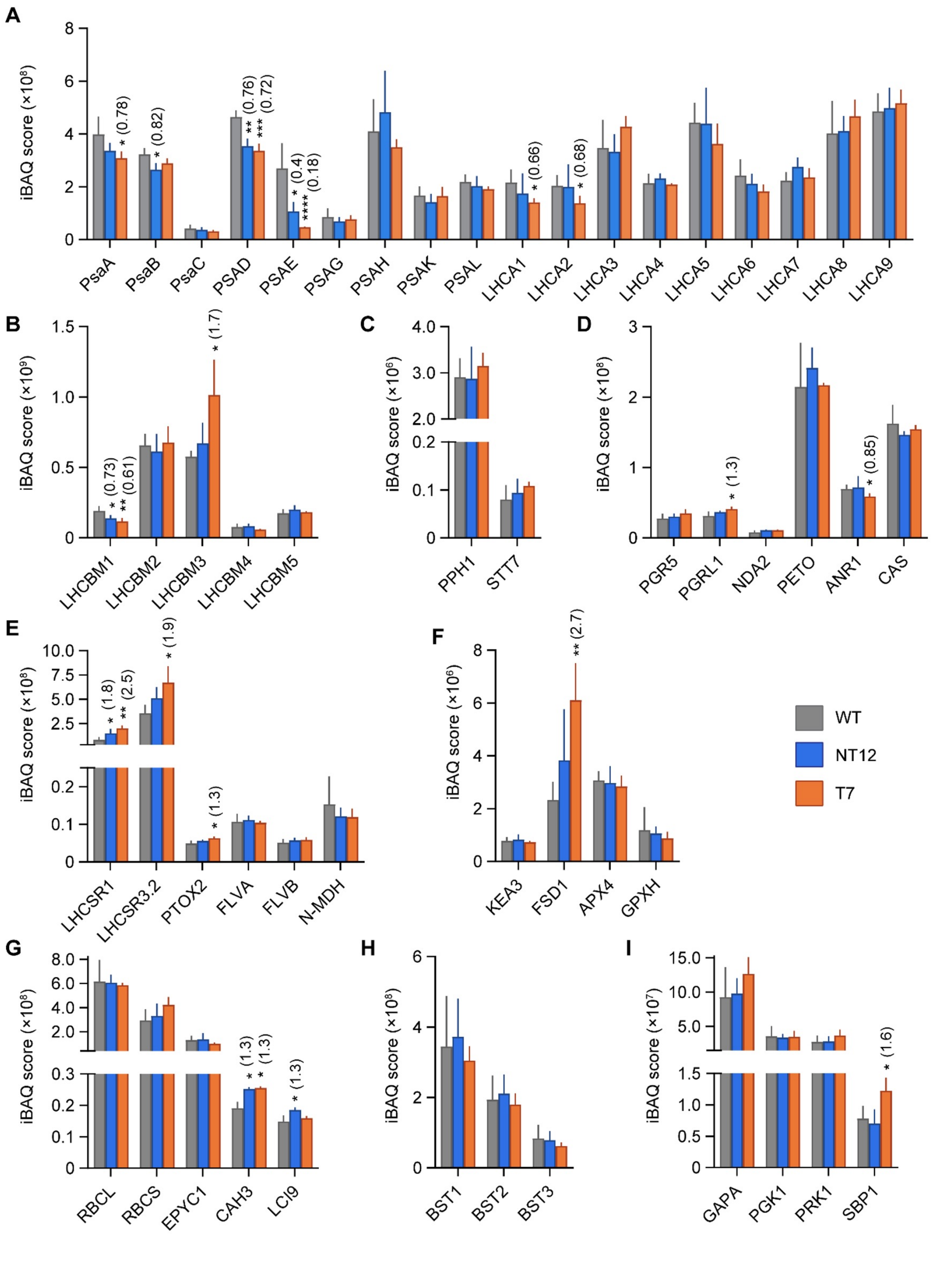


**Supplementary Figure S3 | Additional quantification of proteins expressed by WT, NT12 and T7 cells.** Mass spectrometry-based label-free quantification of selected photosynthetic proteins. Bars show means ± SDs. Means in panels A-F are the same as those plotted in the heat maps in Fig 4. **A**, PSI subunits and light harvesting complex I (LHCI) proteins. **B,C**, State transition associated proteins. **D**, Cyclic electron transfer (CET) associated proteins. **E,F**, Stress associated proteins. **G**, Rubisco large (RBCL) and small (RbcS1-2) subunits, essential pyrenoid component 1 (EPYC1), carbonic anhydrase 3 (CAH3) and low $CO_2$-inducible protein (LCI9). **H**, Bestrophins 1-3 (BST1-3; quantified by the sum of isoform-specific peptide ion intensities). **I**, Glyceraldehyde-3-phosphate dehydrogenase (GAPA), phosphoglycerate kinase (PGK1), phosphoribulokinase (PRK1) and sedoheptulose-1,7-bisphosphatase (SBP1). Note that the y-axes scales vary between panels. *Q* values derived from modified t-tests incorporating permutation-based false discovery rate with 250 randomizations ($n = 4$) are shown as 0.01-0.05 (*) and 0.001-0.01 (**), 0.0001-0.001 (***) and < 0.0001 (****), with exact *q* values listed in Source data. Fold change relative to WT is shown in parentheses for differences with $q \leq 0.05$; differences with $q \geq 0.05$ (not significant) are not shown. Supports Fig. 4.

### Supplemental Fig 4

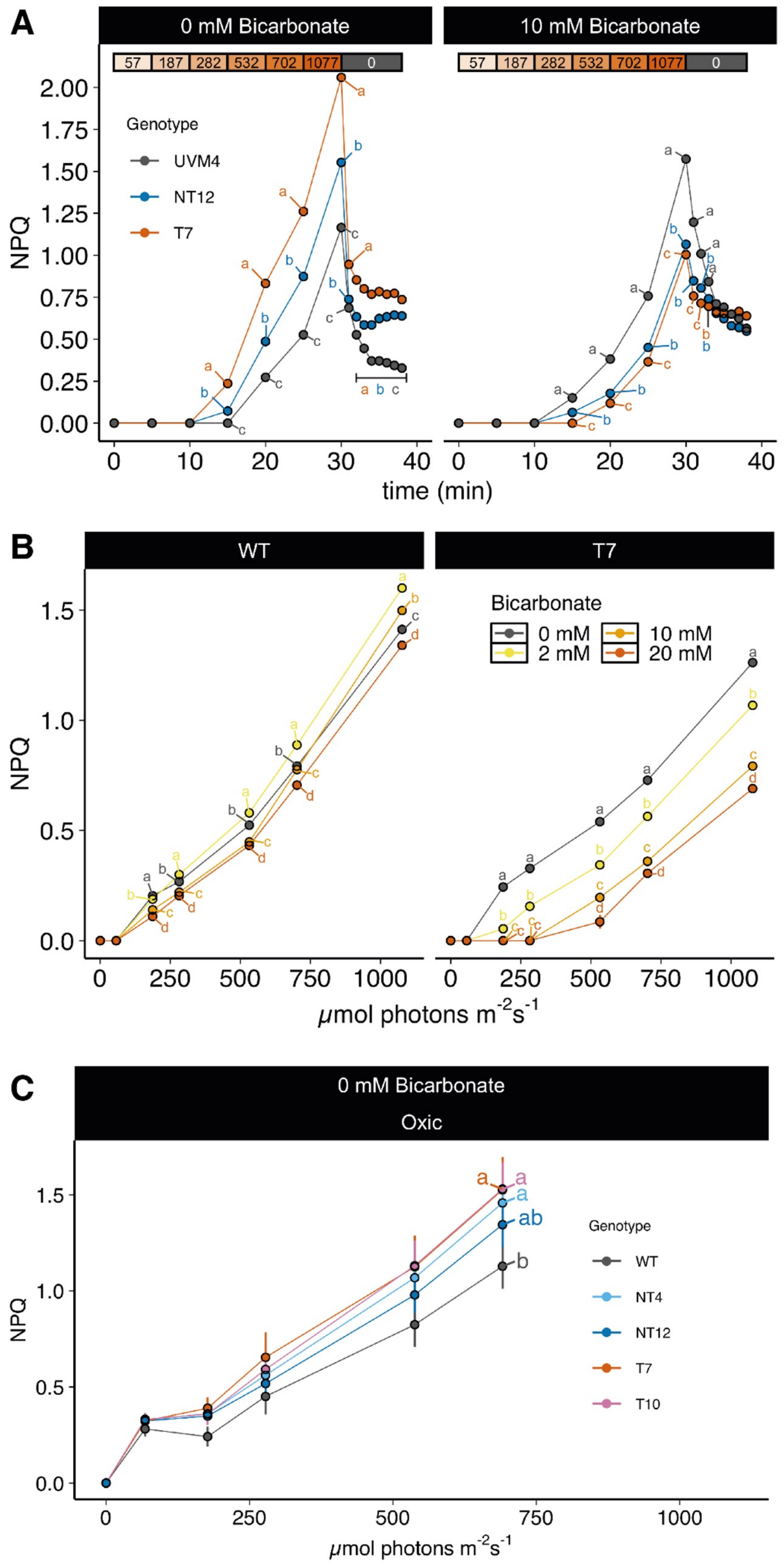


**Supplementary Figure S4 | Effect of added sodium bicarbonate on NPQ in oxic cells. A,** Effect of 0 or 10 mM added sodium bicarbonate on NPQ recorded at different light intensities (shown in orange bars) over a 30-minute illumination. Relaxation of NPQ in the subsequent dark period. **B**, Light curve of NPQ versus light intensity at different concentrations of added sodium bicarbonate. **C**, Comparison of NPQ of additional T and NT mutants and WT oxic cells without additional sodium bicarbonate. Means were compared between genotypes using an ordinary two-way ANOVA and corrected using Tukey's multiple comparison test followed by a HSD test with alpha = 0.05. Different letters indicate significant differences between data points. Supports Fig. 5.

### Supplemental Fig 5

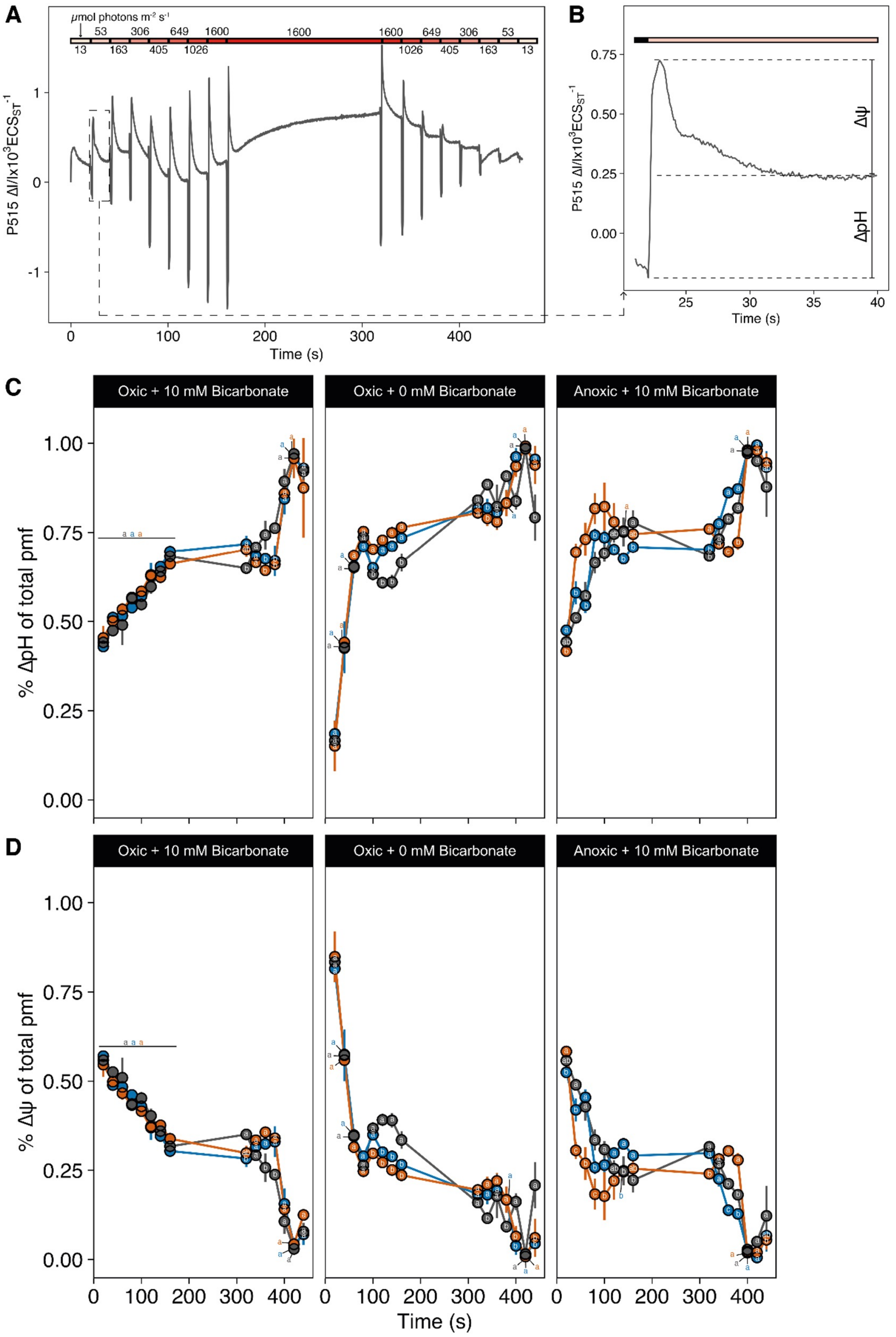


**Supplementary Figure S5 | Partitioning of proton motive force into ΔpH and Δψ. A**, Example raw trace of the P515 signal of Chlamydomonas cells. Light intensity was increased every 20 s (shown at the top), followed by a 1.4 s dark pulse. Cells were then illuminated at the highest light intensity for 10 min, followed by a decrease every 20 s. **B**, Illustration of how pmf was partitioned into ΔpH and Δψ. **C**, Percent ΔpH of total pmf under three different conditions. **D**, Percent Δψ of total pmf under three different conditions. Colours indicate different genotypes. Means were compared between genotypes using an ordinary two-way ANOVA and corrected using Tukey's multiple comparison test followed by a HSD test with alpha = 0.05. Different letters indicate significant differences between data points. Supports Fig. 6.

### Supplemental Fig 6

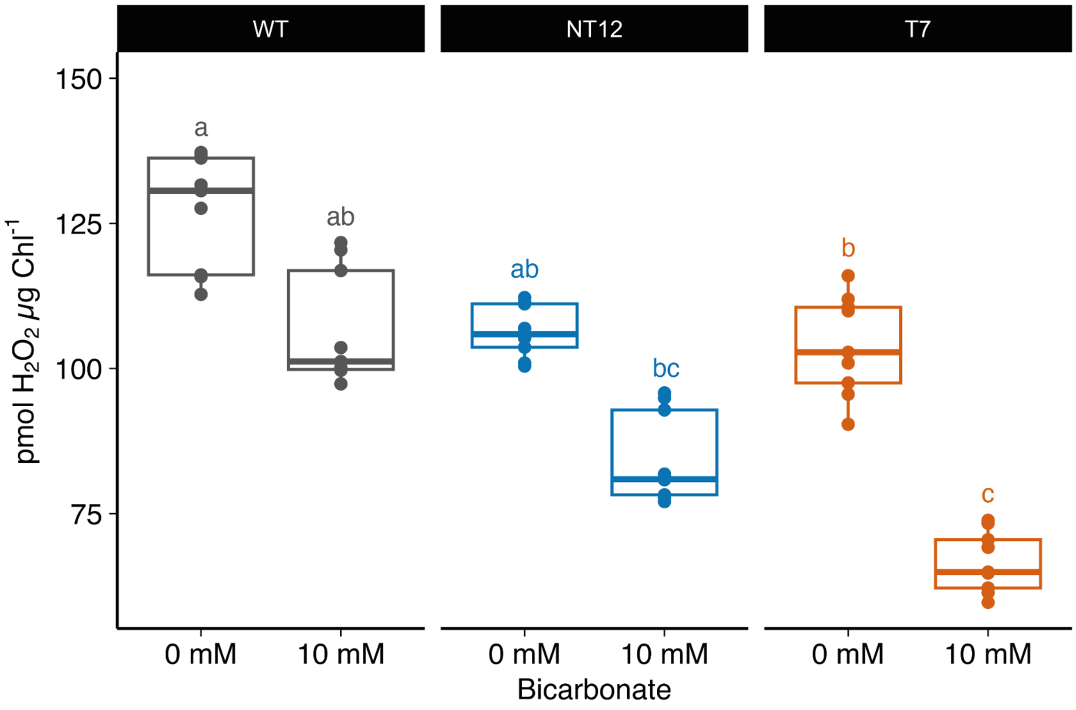


**Supplementary Figure S6 | Hydrogen peroxide levels in NT12, T7 and WT cells.** Cells were grown in TAP medium and switched to TP for at least 24 h. 1 mL of culture was diluted with 1 mL fresh TP containing 1 U of horseradish peroxidase and 5 µM of Amplex Red and incubated under red light at 520 µmol photons $m^{-2}$ $s^{-1}$ for 1 h. $H_2O_2$ in the supernatant was subsequently quantified using a linear $H_2O_2$ standard. Box lines represent the median and first and third quartiles; whiskers indicate the range; circles indicate the raw data. Means were compared between genotypes using an ordinary two-way ANOVA and corrected using Tukey's multiple comparison test followed by a HSD test with alpha = 0.05. Different letters indicate significant differences between genotypes and $CO_2$ concentrations. Supports Fig. 7.

### Supplemental Fig 7

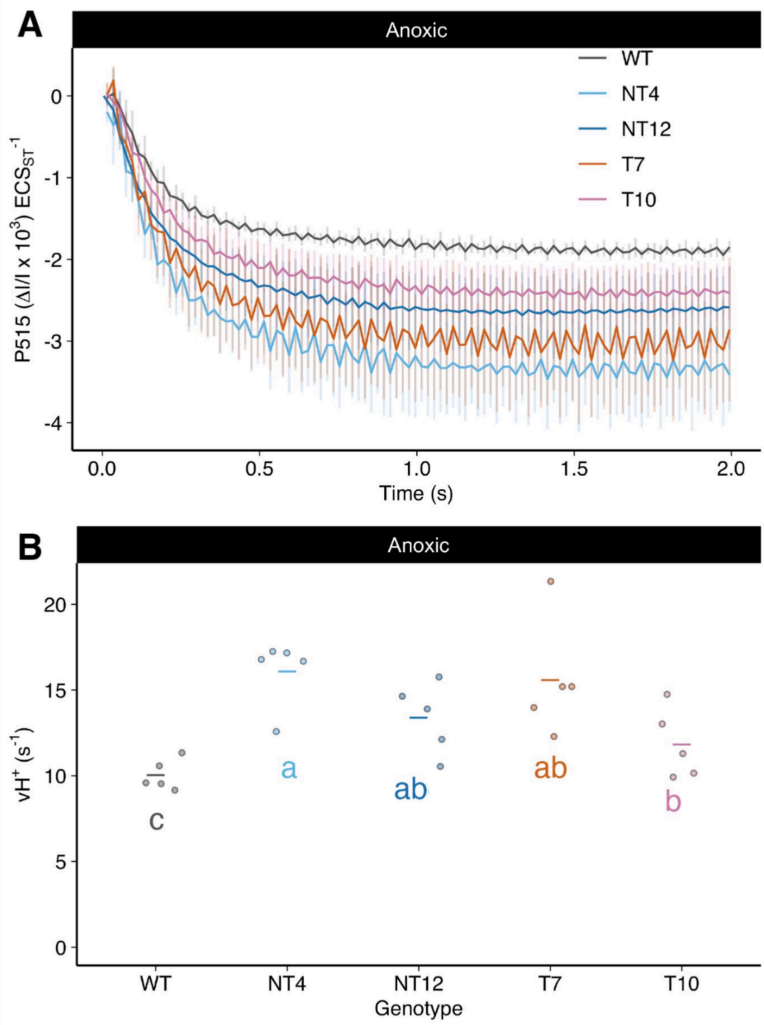


**Supplementary Figure S7 | Rate of ECS absorption decay under CET conditions in T and NT mutant lines relative to WT. A**, Initial rate of ECS decay in WT, NT4, NT12, T7 and T10 treated with DCMU/HA under anoxic conditions upon cessation of a 10 second illumination period using 502 µmol photons $m^{-2}$ $s^{-1}$. **B**, Calculated initial rate ($vH^+$) of ECS decay under the same conditions. Horizontal lines represent means and symbols raw data points. Means were compared between genotypes using an ordinary two-way ANOVA and corrected using Tukey's multiple comparison test followed by a HSD test with alpha = 0.05. Different letters indicate significant differences between data genotypes. Supports Fig. 9.

### Supplemental Fig 8

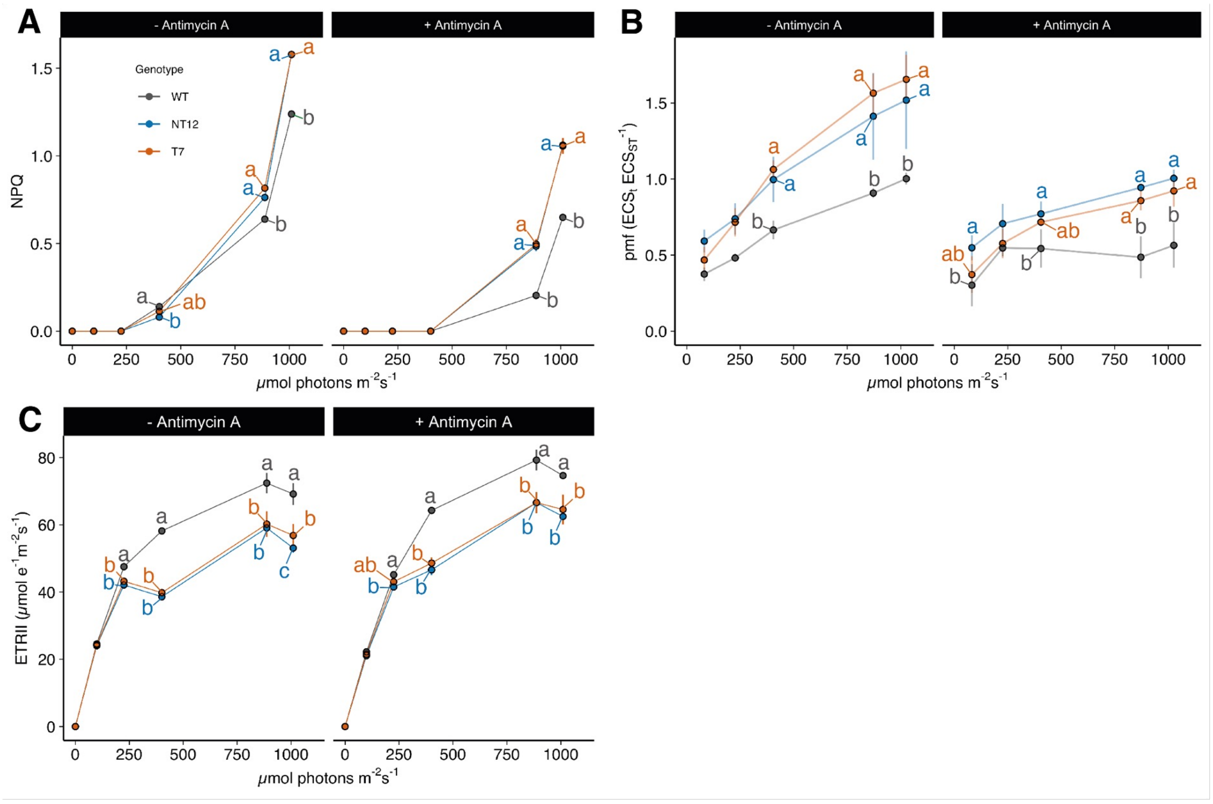


**Supplementary Figure S8 | Effect of Antimycin A on chlorophyll fluorescence and ECS parameters in WT, NT12 and T7 cells under anoxia with 10 mM $NaHCO_3$. A**, non-photochemical quenching (NPQ) ± Antimycin A (AA). **B**, proton motive force (pmf) ± AA. **C**, Electron transport rate (ETRII) ± AA. Measurements were made as described in Figs. 5 and 6 except that 40 µM Antimycin A from a 40 mM ethanolic stock was added prior to measurements. Error bars are S.D. (at least replicates per strain). Means were compared between genotypes using an ordinary two-way ANOVA and corrected using Tukey's multiple comparison test followed by a HSD test with alpha = 0.05. Different letters indicate significant differences between data points at each light intensity. Supports Fig. 9.
